# Dynamics of X chromosome hyper-expression and inactivation in male tissues during stick insect development

**DOI:** 10.1101/2024.07.01.601468

**Authors:** Jelisaveta Djordjevic, Patrick Tran Van, William Toubiana, Marjorie Labédan, Zoé Dumas, Jean-Marc Aury, Corinne Cruaud, Benjamin Istace, Karine Labadie, Benjamin Noel, Darren J Parker, Tanja Schwander

## Abstract

Differentiated sex chromosomes are frequently associated with major transcriptional changes: the evolution of dosage compensation (DC) to equalize gene expression between the sexes and the establishment of meiotic sex chromosome inactivation (MSCI). Our study investigates the mechanisms and developmental dynamics of dosage compensation and meiotic sex chromosome inactivation in the stick insect species *T. poppense*. Stick insects are characterized by XX/XO sex determination and an X chromosome which likely evolved prior to the diversification of insects over 450 Mya. We generated a chromosome-level genome assembly and analyzed gene expression from various tissues (brain, gut, antennae, leg, and reproductive tract) across developmental stages in both sexes. Our results show that complete dosage compensation is maintained in male somatic tissues throughout development, mediated by upregulation of the single X chromosome. Contrarily, in male reproductive tissues, dosage compensation is present only in the early nymphal stages. As males reach the 4th nymphal stage and adulthood, X-linked gene expression diminishes, coinciding with the onset of MSCI. This reduction is associated with histone modifications indicative of transcriptional silencing, aligning with meiotic progression. These findings reveal the dynamic regulation of X-linked gene expression in *T. poppense*, and suggest that reduced X-expression in insect testes is generally driven by MSCI rather than an absence of dosage compensation mechanisms. Our work provides critical insights into sex chromosome evolution and the complex interplay of dosage compensation and MSCI across tissues and developmental stages.

## Introduction

In species where sex is determined by differentiated sex chromosomes, sex chromosome copy number differs between males and females. For example, in X0 and XY systems, males have only one copy of each gene located on the X chromosome, while females have two. Because gene copy number generally correlates with expression levels [1], the expression level of X-linked genes in males is expected to be half that of the same genes in females. However, selection can act on the regulation of expression in a sex-specific manner, and potentially restore similar expression levels in both sexes [2–4], a process known as dosage compensation [2, 5, 6]. The mechanisms underlying dosage compensation between the sexes vary with respect to whether sex chromosomes are downregulated in the homogametic sex, or upregulated in the heterogametic sex. Thus, downregulation of X-linked expression in females underlies dosage compensation in eutherian mammals and the worm *Caenorhabditis elegans*: In eutherian mammals, females inactivate one of the X chromosomes in each cell [6, 7], whereas in *C. elegans*, the expression of both X chromosome copies in hermaphrodites is reduced to achieve similar total expression levels as for the single X in males [8, 9]. Upregulation of the single X in males appears to underlie dosage compensation in a diversity of insect species [10–13]. The ancestral sex determination system in insects was most likely XO/XX or XY/XX male heterogamety [14], and the ancient insect X chromosome has likely persisted for over 450 MYA as its content is still conserved in many insect orders [15]. Most insights for the molecular mechanisms underlying dosage compensation in insects are however for species with derived sex chromosomes, such as the fruit fly *Drosophila melanogaster* and the mosquito *Anopheles gambiae*. These two species have a homologous X chromosome which is independently derived from the same autosome during the diversification of dipterans [16]. In the fly, upregulation of the single X in male somatic tissues is partially mediated by the male-specific lethal (MSL) protein complex, which promotes the transcription of X-linked genes via H4K16 hyperacetylation, with apparently no chromosome-wide modification of X expression in females [17, 18]. In the mosquito, dosage compensation is regulated via sex-specific alternative splicing of a gene referred to as “*sex chromosome activation (SOA)”* or *“007”*, but how this gene activates expression on the male X remains unknown [19, 20].

Interestingly, in all studied XO and XY species that feature dosage compensation via X upregulation in males, the single X in males is much less expressed in the testes than in the somatic tissues [11–13, 21]. This is thought to be due to the presence of dosage compensation in somatic tissues and the absence of dosage compensation in the reproductive tissues, perhaps because the gene expression optima in male versus female gonads are so different that there is no or reduced selection for dosage compensation in this tissue [11]. Furthermore, transcriptional silencing of the X because of meiotic sex chromosome inactivation (MSCI) can specifically reduce the expression levels of X-linked genes in the testes of some species [22, 23].

Beyond the specific case of testes in adult males, the extent of dosage compensation variation across tissues and development remains understudied [21, 24]. Furthermore, the key mechanisms affecting X-linked expression in males (dosage compensation and meiotic sex chromosome inactivation) are only characterized for a few well-studied model organisms as mentioned above, preventing inferences of conserved versus lineage-specific aspects of sex chromosome expression. Here we fill these knowledge gaps by exploring how the extent of dosage compensation varies over development in somatic and reproductive tissues, through which mechanisms dosage compensation is achieved, as well as the potential impact of meiotic sex chromosome inactivation in the stick insect *Timema poppense*. These stick insects are characterized by XX:XO sex determination [25] and the ancient insect X chromosome, which likely persisted for more than 450 million years [15]. Previous work on *Timema* stick insects showed that in adults, there is dosage compensation in the head and legs, but not in the reproductive tract [13]. We generate a new chromosome-level genome assembly of *T. poppense* based on long-read sequencing and Hi-C based scaffolding, and sequence RNA from brain, gut, antennae, leg and reproductive tract samples in males and females across development. We then assess the expression levels of genes located on the X chromosome(s) in both sexes relative to those on the autosomes. Finally, we test for meiotic sex chromosome silencing in the male germline via immunolabeling and genome profiling for silencing histone marks, and discuss how MSCI globally affects X-linked gene expression patterns in male gonads.

## Material and methods

### Sample collection and reference genome generation

We used wild-collected females of the species *T. poppense* (36°59’44.5“N 121°43’05.0“W) for generating the reference genome. We assembled contigs based on Nanopore and Illumina libraries (sequenced at 75x respectively 50x coverage) generated from a single female, and then scaffolded contigs using a Hi-C library (sequenced at 61x coverage) based on a different female. We annotated this assembly using transcriptome data from different tissues and development stages of male and female *T. poppense* as well as from a closely related species (*T. douglasi*; see Gene annotation section).

### Sequencing libraries

To extract high molecular weight (HMW) DNA, we flash-froze a single female (without gut) in liquid nitrogen and ground it using a Cryomill (Retsch). We then extracted HMW DNA using a G/20 Genomic Tips kit (Qiagen) following the manufacturer’s protocols. We checked DNA integrity on a pulse field agarose gel.

A total of 7 ONT libraries were prepared following Oxford Nanopore instructions. One library was prepared using the SQK-LSK108 ligation sequencing kit and was loaded on a MinION R9.4.1 Flow Cell, and six libraries were prepared using the SQK-LSK109 ligation sequencing kit and were loaded on two MinION flow cells and 4 PromethION R9.4.1 Flow Cells. Flow Cell loading was performed according to the Oxford Nanopore protocol and resulted in 78X coverage.

A PCR free Illumina library (based on the same HMW extraction) was prepared using the Kapa Hyper Prep Kit (Roche, Basel, Switzerland), following manufacturer instructions. The library was quantified by qPCR using the KAPA Library Quantification Kit for Illumina Libraries (Roche), and the library profile was assessed using a High Sensitivity DNA kit on an Agilent Bioanalyzer (Agilent Technologies, Santa Clara, CA, USA). The library was then sequenced to approximately 50x coverage on an Illumina HiSeq 4000 instrument (Illumina, San Diego, CA, USA), using 150 base-length read chemistry in a paired-end mode.

A Hi-C library was prepared using the Proximo Hi-C Kit. We generated ground tissue for cross-linking using Cryomill (Retsch) and following manufacturer instructions, using a different female than the one used for Nanopore and HiSeq sequencing, from the same natural population. Library construction and sequencing (250 Mio read pairs) was outsourced to Phase Genomics (Seattle).

### Assembly pipeline and parameters

Raw Oxford Nanopore reads were filtered using Filtlong v0.2.0 (https://github.com/rrwick/Filtlong) with the parameters --min_length 1000 --keep_percent 90 --target_bases 69050000000. The filtered Nanopore reads were then assembled into contigs using Flye v2.8.1 [26] with --genome-size 1.3 Gbp. All Nanopore reads were mapped against the contigs using minimap2 v2.19 [27] with the parameters - c -x map-ont and a first step of polishing was performed using Racon v1.4.3 [28]. Three additional rounds of polishing were then conducted using the Illumina short reads. The short reads were aligned to the contigs using BWA mem v0.7.17 [29] and polishing was performed using Pilon v1.23 [30].

The assembly was decontaminated using BlobTools v1.0 [31] under the taxrule “bestsumorder”. Hit files were generated after a blastn v2.10.1+ against the NBCI nt database, searching for hits with an e-value below 1e-25 (parameters: -max_target_seqs 10 -max_hsps 1 -evalue 1e-25). Contigs without hits to metazoans were removed. Haplotypic duplications were filtered out: filtered reads were mapped against the decontaminated genome using minimap2 and haplotigs were detected with Purge Haplotigs v1.1.1 [32] using the parameters -l 3 -m 17 -h 190 -j 101 following the recommendations by [32].

For scaffolding, Hi-C reads were mapped to the haploid genome using Juicer v1.6 [33] with the restriction site Sau3AI. Chromosome-level scaffolding was then performed using 3D-DNA v180922 [34] with the parameters --editor-coarse-resolution 25000, as recommended by the authors. The resulting Hi-C contact matrices were visualized with Juicebox, and polished following the recommendations by [33]. The completeness of the assembly was assessed with BUSCO v5.1.2 [35] and the insecta_odb10 dataset using the --long and --augustus parameters.

To identify the X chromosome in our assembly, we used a coverage approach. We compared coverage between males and females because *Timema* have XX/X0 sex determination [25] and males are expected to show half of the female coverage at the X chromosome. We mapped 3 female (SRS7637462, SRS7637490, and SRS7638308 from [36]) and 4 male samples (PRJNA725673 from [13]) to our scaffolded genome which allowed us to unambiguously identify the third largest scaffold as the X chromosome (Supplemental figure 1).

The *T. poppense* genome was annotated using a combination of *ab initio* gene prediction, protein homology, and RNA-seq using the Braker2 pipeline v. 2.1.6 [37]. To begin, the genome assembly was soft-masked using RepeatModeler (v. 2.0.2, options: -LTRStruct, -engine ncbi) and RepeatMasker (v. 4.1.2, options: -engine ncbi, -xsmall). For protein evidence, we used the arthropod protein sequences from OrthoDB v.10.1 [38] and the predicted protein sequences for *Timema* from our previous genome assemblies [36]. For RNAseq evidence, we used our newly generated RNAseq data for *T. poppense* (175 libraries, see below) as well as publicly available RNAseq data from *T. poppense* and the closely related species *T. douglasi* (Bioproject Accessions: PRJNA380865, PRJNA679785, PRJNA1128519) for a total of 376 RNAseq libraries (364 paired-end, 12 single-end) covering 117 different life stages, tissue, and sex combinations. Reads were quality trimmed with Trimmomatic (v. 0.39, options: ILLUMINACLIP:3:25:6 LEADING:9 TRAILING:9 SLIDINGWINDOW:4:15 MINLEN:80) [39] before mapping to the genome assembly with STAR (v. 2.7.8a, options: --twopassMode Basic) [40]. Braker2 was run using protein evidence and RNAseq separately with the gene predictors Augustus v. 3.4.0 [41] and Genemark v. 4.72 [42]. Following the RNAseq run, UTR predictions were added to the RNAseq gene predictions using GUSHR v. 1.0 [43] in Braker2 (--addUTR=on). The separate gene predictions were then merged using TSEBRA v1.0.3 [44] using the pref_braker1.cfg configuration file, which weights RNAseq evidence more strongly than the default option. We then ran BUSCO (v. 5.3.2, insecta_odb10) on the gene regions annotated by Braker2 and on the whole genome assembly. Any genes found by BUSCO but missed by Braker were then added to the annotation (48 genes). ncRNA genes were predicted using Infernal (v. 1.1.2, minimum e-value 1e-10) [45]. GO-terms for protein-coding gene predictions were obtained using blastP within OmicsBox v3.1.2, default parameters) to blast the nr *Drosophila melanogaster* database (taxonomy filter: 7227).

### Insect husbandry

Hatchlings were obtained from eggs laid by captive bred *T. poppense* individuals, originally collected in California in 2018. To complete development, *Timema* females have one extra molt compared to males [46], hence our aim was to obtain five different tissues (reproductive tracts and four somatic tissues) for each of seven developmental stages in females (nymphal stage 1-6, adult) and six in males (nymphal stage 1-5, adult; Supplemental Table S1). Upon hatching, insects were reared in Petri dishes containing *Ceanothus* plant cuttings wrapped in wet cotton up to a specific developmental stage and then dissected one day after molting. To identify individuals that molted, we painted the thoraxes with red acrylic paint after each molt and checked daily for individuals without the paint. Prior to dissection, the insects were anesthetized with CO_2_. Brain, antennae, legs, and gut samples were obtained from each developmental stage, while the reproductive tract was collected at every stage from the 4th nymphal stage (Supplemental Table S1). We dissected reproductive tracts only starting from the 4th nymphal stage because we were originally unable to unambiguously identify gonad tissues in the earlier stages. Upon improving our dissection techniques, we were later able to identify and dissect gonad tissues at earlier stages, and we therefore secondarily added reproductive tract samples of newly hatched individuals (1^st^ nymphal stage). During dissections, tissues were placed in Eppendorf tubes and immediately flash frozen in liquid nitrogen before storage at −80°C until extraction for approximately 1/3 of the dissections. For the remaining dissections, due to pandemic-related laboratory closures, tissues were preserved in RNA later (Qiagen) before storage at −80°C. In total, we obtained two to four replicates per sex and tissue (Supplemental Table S1).

### RNA extraction and sequencing

TRIzol solution (1 mL) and a small amount of ceramic beads (Sigmund Lindner) were added to each tube containing tissue. Samples were homogenized using a tissue homogenizer (Precellys Evolution; Bertin Technologies). Chloroform (200 μL) was added to each sample and samples were then vortexed for 15 seconds and centrifuged for 25 minutes at 12,000 revolutions per minute (rpm) at 4°C. The upper phase containing the RNA was then transferred to a new 1.5 mL tube with the addition of isopropanol (650 μL) and Glycogen blue (GlycoBlue™ Coprecipitant; 1 μL). The samples were vortexed and placed at −20°C overnight. Samples were then centrifuged for 30 minutes at 12,000 rpm at 4°C. The liquid supernatants were removed, and the RNA pellet underwent two washes using 80% and 70% ethanol. Each wash was followed by a 5-minute centrifugation step at 12,000 rpm. Finally, the RNA pellet was resuspended in nuclease-free water and quantified using a fluorescent RNA-binding dye (QuantiFluor RNA System) and nanodrop (DS-11 FX).

Library preparation using NEBNext (New England BioLabs) and sequencing on an Illumina NovaSeq 6000 platform with 100 bp paired-end sequencing (∼45 million read pairs per sample on average) was outsourced to a sequencing facility (Fasteris, Geneva).

### Data analyses

#### Read counting

Aligned RNA-Seq reads, sorted by genomic position, were processed to count reads mapped to exonic regions using HTseq [47] (v.011.2, options: --order=pos --type=exon --idattr=gene_id --stranded=reverse).

#### Dosage compensation analysis

We categorized the data based on tissue and developmental stage. Subsequently, we excluded genes with low expression, specifically those that were not expressed in a minimum of three libraries when the replicate number equaled four, two libraries when the replicate number was three, and one library when the replicate number was two in each sex. This filtering required an expression level greater than 0.5 CPM (counts per million) in males or females within the specified tissue and stage.

For the computation of RPKM values, we first obtained gene sizes using the GenomicFeatures package [48] and then calculated the average RPKM values for each sex using EdgeR [49]. Lastly, we determined the log2 RPKM ratio between the male and female average expression levels. Statistical analyses (Wilcoxon test), graphical representations of the data were performed in R (4.3.1).

#### Tissue specificity (Tau)

To test if the X chromosome has more tissue-specific expression than the autosomes, we calculated Tau, an index of gene expression tissue specificity ranging from zero (expressed equally in all studied tissues) to one (gene expressed in only one tissue) [50]. For these analyses we used the subset of four tissues (antenna, brain, gut and reproductive tract) and four developmental stages (1^st^, 3^rd^, 4^th^ nymphal and the adult stage) for which we had data for both sexes (see Supplemental Table S1). Using median expression RPKM values, we calculated Tau for the 12 longest scaffolds (corresponding to the 12 chromosomes of *T. poppense*). We visualized the results with ggplot2 (v. 3.3.2) [51].

#### Immunolabeling of X chromosomes

In order to determine the presence of meiotic cells as well as X heterochromatization and transcriptional activity in male gonads, we performed double immunolabeling of SMC3_488 (#AB201542, AbCam) as a marker of cohesin axes (this allows for the assessment of synapsis progression, and hence, cell cycle) and either H3K9me3 (#AB8898, AbCam; which labels silencing heterochromatin marks) or RNA polymerase II phosphorylated at serine 2 (p-RNApol-II (ab193468, AbCam); an indicator of transcription). Gonads from 1^st^ nymphal and 4^th^ nymphal males were dissected in 1X PBS, then fixed in paraformaldehyde (2%) and Triton X-100 (0.1%) solution for 15 minutes. Gonads were then squashed on slides coated with poly-L-lysine, and flash frozen in liquid nitrogen. Slides were incubated in PBS 1x for 20 minutes at room temperature, then blocked with a BSA 3% solution (prepared by dissolving 3g of BSA in 100mL of PBS 1X) for 20 minutes. Slides were then incubated overnight at 4°C in a humid chamber with the primary (p-RNA-Pol II or H3K9me3), diluted in BSA 3% (1:100), and washed 3x for 5 minutes in PBS 1X. The secondary antibody, anti-rabbit Alexa 594 (711-585-152, Jackson), diluted in BSA 3% (1:100), was applied, and samples were incubated for 45 minutes at room temperature. Further washes were conducted as described above, followed by an extended 10-minute wash in PBS 1X, and by blocking using a 5% Normal Rabbit Serum (NRS) solution (X090210-8, Agilent Technology) for 30 minutes at room temperature. Another 5-minute wash in PBS 1x was performed to remove excess blocking solution. Samples were then incubated with the labeled antibody SMC3_488 (#AB201542, AbCam) at a 1:100 dilution for 1 hour at room temperature, followed by a final series of washes and staining with DAPI (D9542, Sigma-Aldrich) for 3 minutes at room temperature. Finally, after a 5-minute wash in 1x PBS, two drops of Vectashield were added to the slide, which was then covered with a coverslip, and sealed with nail varnish for imaging. Image acquisition was performed at the Cellular Imaging Facility (CIF, University of Lausanne) using a Zeiss LSM 880 Airyscan.

#### ChIP-sequencing to reveal heterochromatin marks

To profile silencing histone modifications on the X chromosome in male gonads, we conducted ChIP-sequencing using H3K9me3 (tri-methylation of histone H3 at lysine-9), a histone modification generally associated with heterochromatic silencing, and which marks the silenced X chromosome in male mouse spermatogenesis (Khalil, et al. 2004; Ernst, et al. 2019). We dissected male gonads and immediately froze them in liquid nitrogen. For chromatin preparations, frozen tissues were homogenized by cryogenic grinding (CryoMill; Retsch GmbH) using a specific regimen (2x 60 s, 25 Hz, resting 30 s, 5 Hz), transferred to a 15-ml Falcon tube, and subsequently subjected to rotation at room temperature for 12 minutes in a cross-linking solution composed of 50 mM Hepes (pH 7.9), 1 mM EDTA (pH 8), 0.5 mM EGTA (pH 8), 100 mM NaCl, and 1% formaldehyde. The cross-linking reaction was stopped by pelleting nuclei for 2 min at 2000g, followed by rotation for 10 minutes in a stop solution containing PBS, 125 mM glycine, and 0.01% Triton X-100. Nuclei were then subjected to washing steps in solution A [10 mM Hepes (pH 7.9), 10 mM EDTA (pH 8), 0.5 mM EGTA (pH 8), and 0.25% Triton X-100] and solution B [10 mM Hepes (pH 7.9), 1 mM EDTA (pH 8), 0.5 mM EGTA (pH 8), 0.01% Triton X-100, and 200 mM NaCl]. This was followed by a sonication step in 100 μl of radioimmunoprecipitation assay (RIPA) buffer [10 mM tris-HCl (pH 8), 140 mM NaCl, 1 mM EDTA (pH 8), 1% Triton X-100, 0.1% SDS, 0.1% sodium deoxycholate, and 1× complete protease inhibitor cocktail] in AFA microtubes in a Covaris S220 sonicator for 5 min with a peak incident power of 140 W, a duty cycle of 5%, and 200 cycles per burst. Sonicated chromatin was centrifuged to pellet insoluble material and snap-frozen.

ChIP was carried out using 5 μl of H3K9me3 (#AB8898, AbCam) incubated overnight at 4°C with half of the prepared chromatin sample (10 ul of the same chromatin preparation was used as input control, see below). Protein A Dynabeads (50 μl; Thermo Fisher Scientific, 100-01D and 100-03D) were added for 3 hours at 4°C, and subsequently washed for 10 min each once with RIPA, four times with RIPA with 500 mM NaCl, once in LiCl buffer [10 mM tris-HCl (pH 8), 250 mM LiCl, 1 mM EDTA, 0.5% IGEPAL CA-630, and 0.5% sodium deoxycholate], and twice in TE buffer [10 mM tris-HCl (pH 8) and 1 mM EDTA]. DNAs of the ChIP sample and the input control were then purified by ribonuclease digestion, proteinase K digestion, reversal of cross-links at 65°C for 6 hours, and elution from a QIAGEN MinElute PCR purification column. The purified DNA (ChIP and input) was then processed at the Lausanne Genomic Technologies Facility for library preparation using the NEBNext Ultra II DNA Library Prep Kit for Illumina and 150-bp paired-end sequencing on an Illumina HiSeq 4000.

ChIP and input reads were trimmed using trimmomatic (v0.39). The reads were then mapped to the reference genome using the BWA-MEM algorithm v0.7.17 (-c 1000000000). Chimeric reads were removed using SA:Z tags, and PCR duplicates were eliminated with Picard tools (v2.26.2). Mean coverage was computed for ChIP and input reads within non-overlapping 30 kb windows across all scaffolds using BEDTools (v2.30.0) and normalized by the number of mapped reads in each library.

## Results and Discussion

### Complete dosage compensation across male somatic tissues and developmental stages

*T. poppense* stick insects have 12 chromosomes, comprising one X chromosome (third in size) and 11 autosomes [25]. Our genome assembly fully matches the structure expected from karyotypes, with 1.26 Gbp (94%) of the 1.34 Gbp assembly comprised in the 12 longest scaffolds (hereafter chromosomes) and scaffold 3 corresponding to the X chromosome (supplemental Figure S1). In all four somatic tissues examined (guts, brains, antennae, legs), males feature complete dosage compensation throughout nymphal development as well as at the adult stage. This is revealed by similar ratios of male to female expression for autosomal and X-linked genes (Figure 1A). Additionally, gene expression levels are relatively constant along the single male X (Supplemental Figure S2). This indicates that dosage compensation is achieved through a global mechanism that affects the entire chromosome.

**Figure 1.**
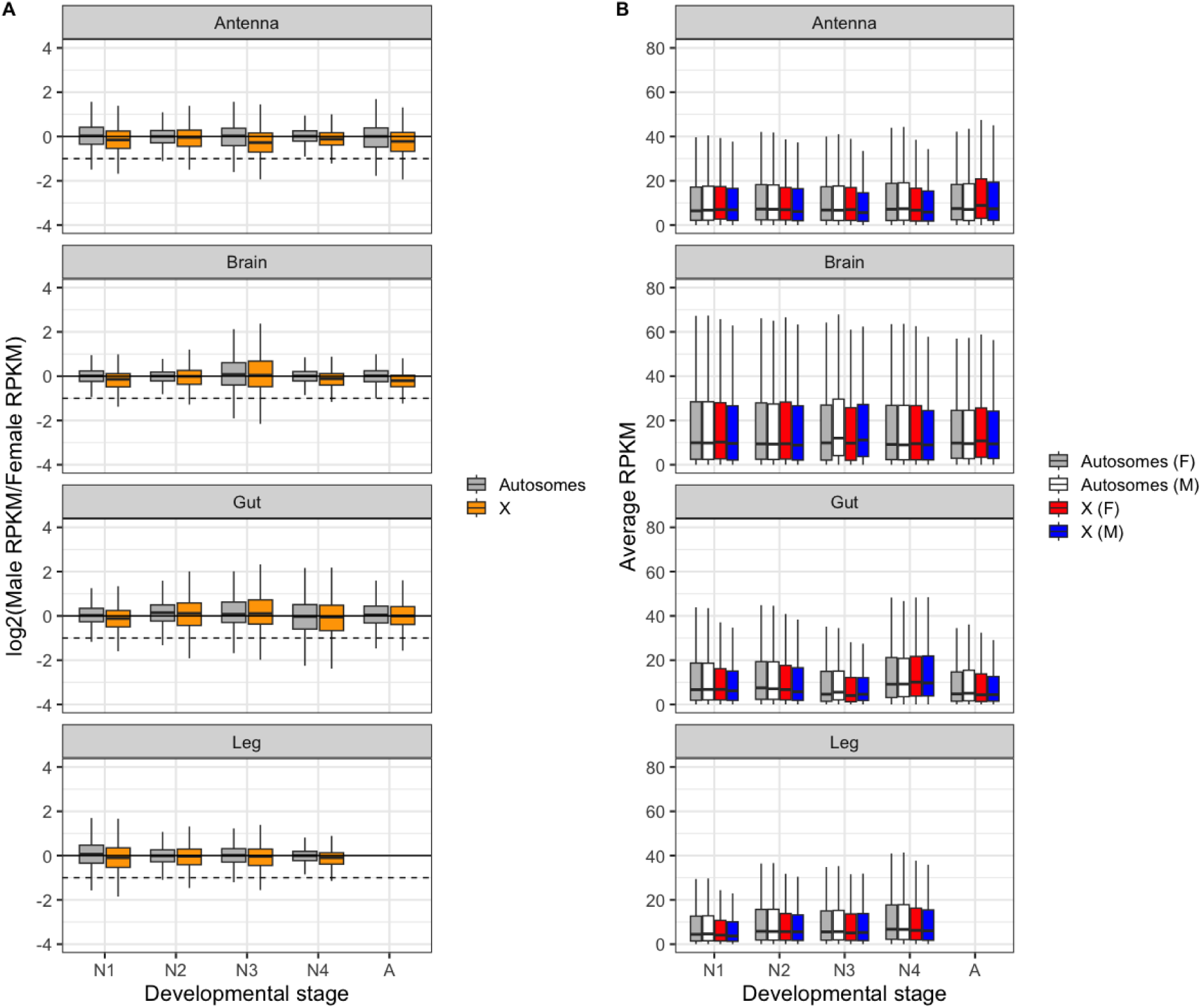
Complete dosage compensation in male somatic tissues during development. **A)** Log2 of male to female expression ratio for the X (orange) and autosomes (grey) in somatic tissues along development. Dashed lines represent a two-fold reduction in expression in males (as expected if there was no dosage compensation). See Supplemental Figure S2 for individual rather than pooled autosomes **B)** Average expression levels (RPKM values) of genes located on the autosomes and X in females (grey and red) and in males (white and blue) in somatic tissues during development. Boxplots depict the median, the lower and upper quartiles, while the whiskers represent the minimum and maximum values, within 1.5x the interquartile range. See Supplemental Figure S3 for individual rather than pooled autosomes.

While dosage compensation is complete between males and females (Figure 1A), X-linked genes generally have significantly lower expression than autosomal genes in male somatic tissues (Figure 1B; Supplemental Table S3). A similar pattern is observed for females (Figure 1B), with a significant reduction of X-linked expression in female gut and gonad tissue; (Supplemental Table S3). It can be argued that complete dosage compensation should refer to situations where the single X is transcribed at a level comparable to the ancestral levels for two X copies, prior to X chromosome formation (i.e., when the X chromosome would have been an autosome; [52]). However, the X chromosome in stick insects has been conserved for at least 120 Mya [13, 53] and shares most of its content with the X chromosome in other insect orders that diverged over 450 million years ago [15], making it impossible to infer ancestral transcription levels.

Why X-linked genes are overall less expressed than autosomal genes in *Timema* remains an open question. A likely explanation is that there is an upper limit to how much the transcriptional activity of a haploid X can be increased, which would favor the movement of highly expressed genes from the X to autosomes [54, 55]. A limit to the maximal transcriptional activity of a haploid chromosome is notably believed to explain X chromosome gene content and reduced expression levels in therian mammals [54, 55] and *Drosophila* fruit flies [56]. Support for a similar explanation in *Timema* stems from the tissue-specificity of X-linked genes. Along with lower levels of expression, the X chromosome in mammals has higher tissue specificity than autosomes [55], because highly expressed genes generally have broader expression [55, 57]. Similar to mammals, we find on average higher tissue specificity of X-linked than autosomal genes in both male and female *Timema* (Wilcoxon test, *p*_adj (females)_= 3.5e-11, *p*_adj (males)_= 2.3e-08; Figure 2; see also Supplemental Figure S4 and Supplemental Table S5).

**Figure 2.**
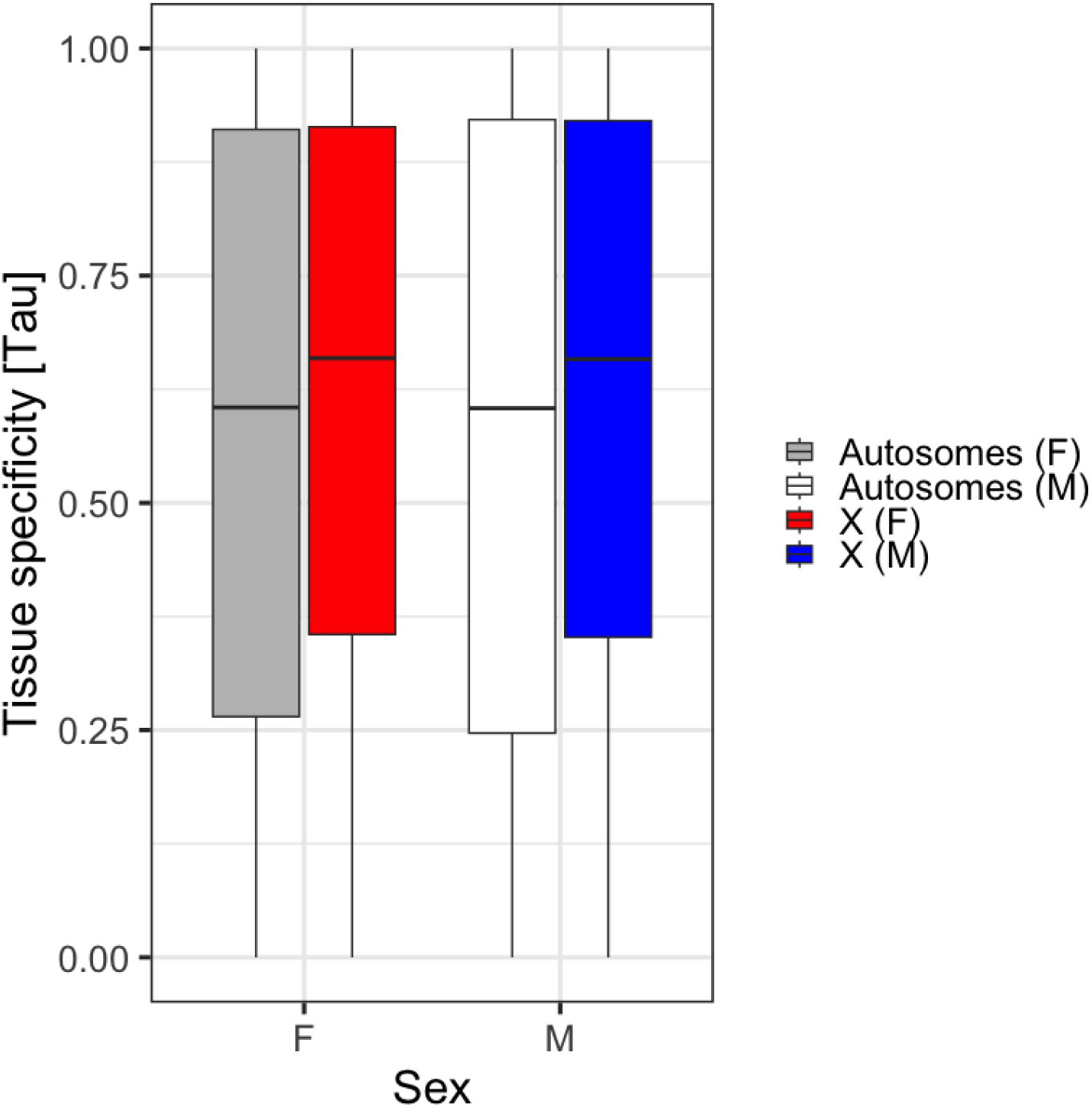
X-linked genes are more tissue-specific in their expression than autosomal genes. Tissue specificity of expression for genes on the X chromosome in males (blue box) and females (red box), compared to autosomal genes (white and gray boxes), calculated across four tissues. Boxes depict the median, the lower and upper quartiles, while the whiskers represent the minimum and maximum values, within 1.5x the interquartile range. Note that this pattern is not solely driven by averaging values across 11 autosomes or gene expression in gonads (Supplemental Figure S4).

Lastly, X chromosome degradation could also contribute to lower expression of X-linked as compared to autosomal genes, similar to patterns reported for genes on the Y chromosomes in some species [58]. The X chromosome has a reduced effective population size as compared to autosomes, as a consequence of reduced copy numbers and the lack of X recombination in males, which leads to the accumulation of deleterious mutations in *Timema* [13].

### Early presence and later absence of dosage compensation in the male reproductive tract

The patterns of X-linked expression in *Timema* male reproductive tracts differ strikingly from those observed in somatic tissues. During the first nymphal stage, there is complete dosage compensation in the reproductive tract, similar to somatic tissues (Figure 3A, B, C). However, from the 4^th^ nymphal stage, there is a strong reduction of X-linked expression in males, relatively constant along the length of the X (Supplemental Figure S6), and which persists throughout development and until the adult stage (Figure 3). We first hypothesized that dosage compensation might be active during early gonad development in the first nymphal stage because the tissue was not yet strongly sexually differentiated. However, this is not the case as the degree of gonadal sexual differentiation (as measured from sex biased gene expression) is relatively constant throughout development. Already 44% of the expressed genes in gonads show sex-biased expression at the first nymphal stage, as compared to 45% at the 4^th^ nymphal stage or 53% in adults (Supplemental Table S4).

**Figure 3.**
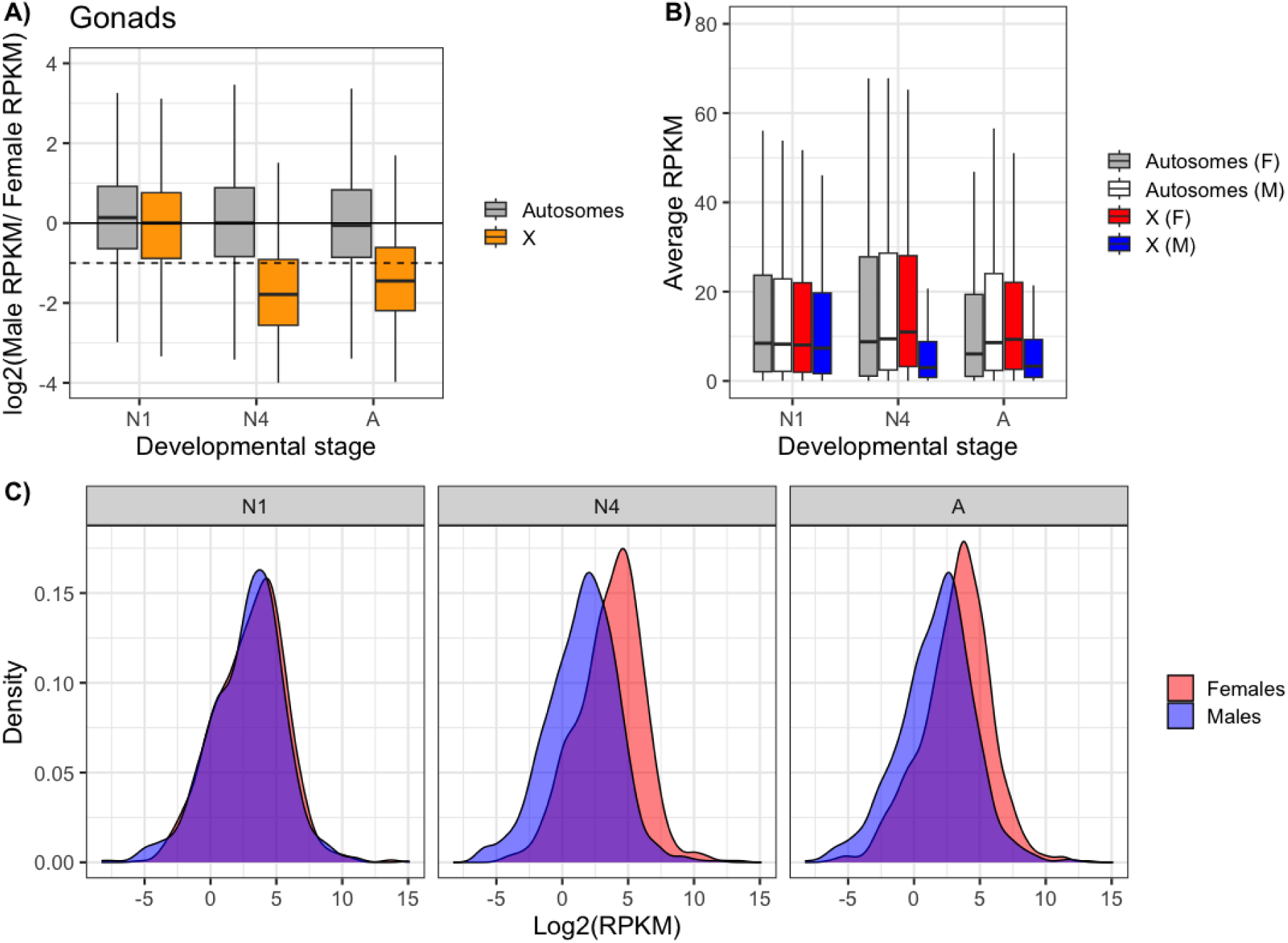
Absence of dosage compensation in male reproductive tracts at 4^th^ nymphal and adult stages. **A)** Log_2_ of male to female expression ratio for the X chromosome (orange) and autosomes (gray) at three developmental stages (N1-1^st^ nymphal, N4-4^th^ nymphal and A-adult stage) in the reproductive tract. See Supplemental Figure S5 for individual rather than pooled autosomes. **B)** Average expression levels (RPKM values) of genes located on the autosomes and X in females (gray and red boxes) and in males (white and blue) at three developmental stages. Boxplots depict the median, the lower and upper quartiles, while the whiskers represent the minimum and maximum values, within 1.5x the interquartile range. See Supplemental Figure S5 for individual rather than pooled autosomes. **C)** Distribution of expression levels for X-linked genes in males (blue) and females (red) (overlapping ranges are indicated in purple).

Next, we evaluated whether the strong reduction of X-linked expression from the 4^th^ nymphal stage was caused by meiotic X-chromosome inactivation, as is well described in therian mammals [59]. Indeed, the extent of the reduction significantly exceeds the two-fold reduction expected in the absence of dosage compensation (Wilcoxon signed rank test with continuity correction; 4^th^ nymphal stage: V = 77366, p-value < 2.2e-16, adult stage: V = 150046, p-value < 2.2e-16) indicating other mechanisms are affecting the expression of X linked genes in male gonads (Figure 3A, B). Immunolabeling of gonad tissue from adult and 4^th^ nymphal stage males corroborates the transcriptional inactivity of the X chromosome. The X chromosome in male meiotic cells is visible in DAPI staining as highly condensed (“heteropycnotic”; Figure 4), similar to other insects [60]. It also lacks a RNApol-II signal, a marker for transcriptional activity, in otherwise transcriptionally active cells (Figure 4). We then also performed immunolabeling of meiotic cells for silencing histone modifications (using an H3K9me3 antibody) and profiled the corresponding molecular marks in the genome using ChIP-seq based on male gonad tissue. As expected, the X has a strong H3K9me3 signal (Figure 5 A, B) and the corresponding H3K9me3 marks are enriched all along the male X (Figure 5 C), suggesting that the silencing of the X is achieved via histone modifications (Figure 5).

**Figure 4.**
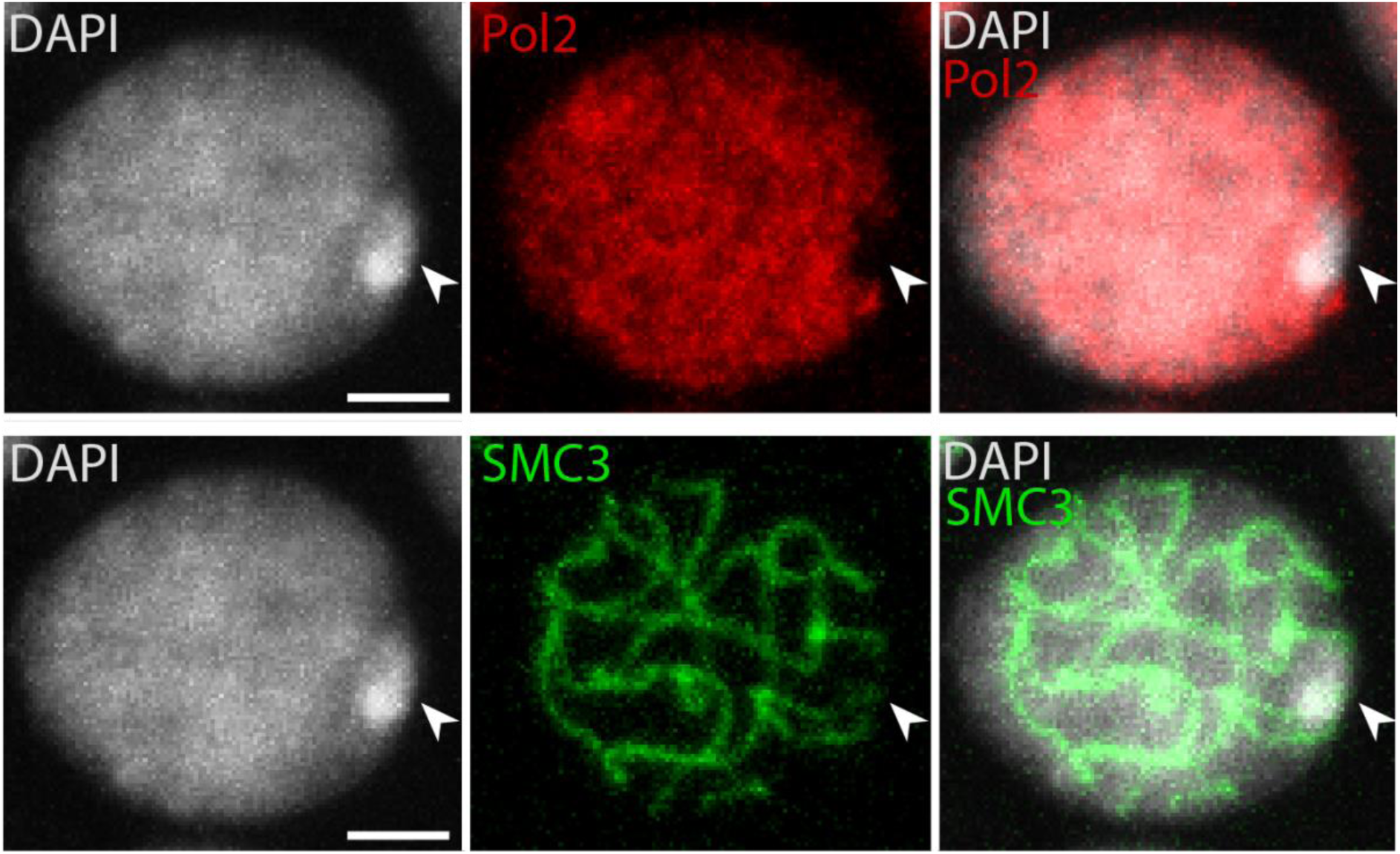
No transcriptional activity of the X chromosome during male meiosis. DAPI, p-RNA Pol-II and SMC3 stainings applied to germ cell squashes of 4^th^ nymphal stage males. Shown is a cell at the pachytene stage of meiosis I for illustration. The arrow in each image indicates the location of the X chromosome. Scale bar indicates 5μm

**Figure 5.**
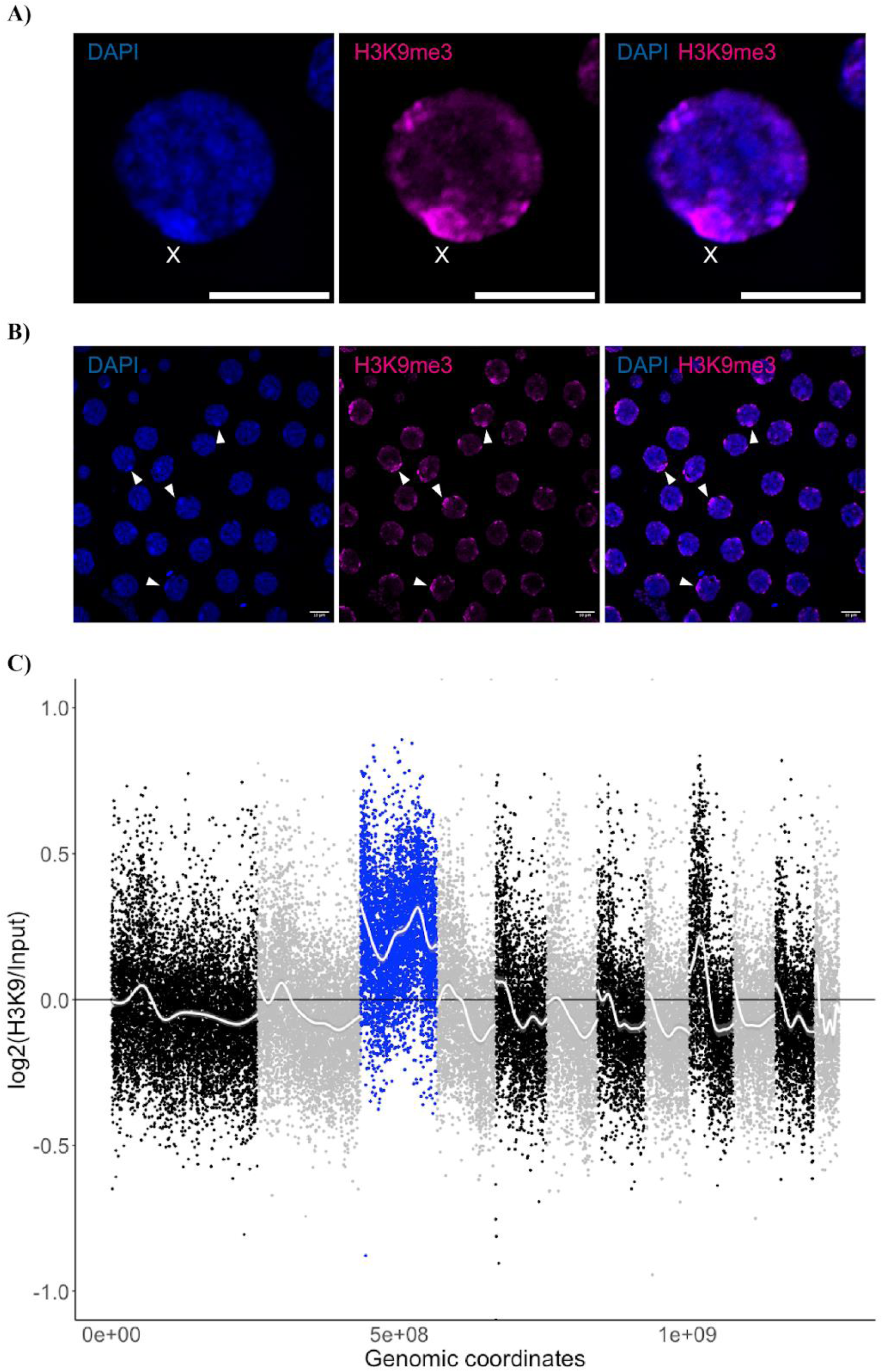
Enrichment of the H3K9me marks on the X chromosome in adult male testes. **A**, **B**) DAPI (blue) and H3K9me3 (magenta) applied to cells from male adult gonad squashes, **A**) single germ cell, with X indicating the location of the X chromosome, scale bar (10µm) **B**) Many germ cells, with arrows pointing to the X chromosome **C**) Log2 ratio of H3K9me to input ChIP-Seq data along the genome, 30K window size. The X chromosome is depicted in blue, while autosomes are depicted in black or gray. Smoothing lines (method “gam”) for each scaffold are in white.

The patterns of MSCI in stick insects are in stark contrast to what is known about X chromosome expression in *D. melanogaster* testis. Indeed, although initially debated, recent evidence indicates that MSCI does not occur in *D. melanogaster*. This is revealed by transcriptional profiling of *D. melanogaster* testis [61–63], similar RNA polymerase activity on the X and autosomes [64], and lack of enrichment of silencing histone modifications on the X in the male germline [64]. As such, the regulation of X chromosome expression in stick insect testes, including MSCI, shares many parallels with mammals and birds [65] but differs strikingly from the more closely related fly species.

At least three, mutually non-exclusive hypotheses have been proposed for the evolution of MSCI. Under the first hypothesis, inactivation would be linked to the lack of synapsis and might have evolved to protect un-synapsed chromosomes from damaging effects such as ectopic exchanges, non-homologous recombination, and unrepaired double-strand breaks [66]. The second hypothesis suggests that MSCI is advantageous because it prevents meiotic expression of selfish genetic elements, including sex-ratio distorters [67, 68]. Finally, the third hypothesis proposes that MSCI is driven by sexually antagonistic effects of X-linked genes [69]. Because the X spends more time in females than in males, it can accumulate female-beneficial genes with detrimental effects in males [70]. It might then be advantageous to reduce or eliminate X-expression in the late stages of spermatogenesis [69].

The presence of MSCI in stick insects as we document here and lack thereof in *D. melanogaster* supports the first hypothesis. A key difference between male meiosis in *Timema* stick insects and *D. melanogaster* is that meiosis is achiasmatic in *Drosophila* males [71] but chiasmatic in *Timema* [13]. The evolution of a completely achiasmatic meiosis in males means that all chromosomes, not only hemizygous sex chromosomes, would be exposed to damaging effects in the absence of synapsis. Such a situation should generate strong selection for sheltering mechanisms not linked to hemizygosity [66].

An association between achiasmy and lack of sex chromosome inactivation is supported by a survey of older cytogenetic data in insects [66]. Chiasmatic insects, including most species in the orders Orthoptera, Blattodea, Mantodea and the suborder Heteroptera, typically show sex chromosome morphologies during male meiosis that likely reflect heterochromatization (i.e., MSCI), while many (though not all) achiasmatic insects do not. This pattern is consistent with the idea that achiasmatic insects evolved from ancestral, chiasmatic forms with sex chromosome inactivation; some achiasmatic groups have subsequently lost sex chromosome inactivation while others have retained it [66].

Independently of the causes underlying the evolution of MSCI, the contrasting X-expression patterns in reproductive tracts in male hatchlings versus 4^th^ nymphal stage likely reflect the start of meiosis I. Thus, in hatchlings where dosage compensation is complete (Figure 3A), cell divisions would be largely mitotic (spermatogonia stage). MSCI would start in cells entering meiosis I, resulting in the X-linked expression pattern we observe in reproductive tracts at the 4^th^ nymphal stage (Figure 3A). In accordance with these interpretations, observation of cells from 4th nymphal stage testes reveal the presence of many cells at meiosis I, whereas we could not find a single cell in meiosis I in squashes from testes of male hatchlings (Supplemental Figure S7). Finally, it appears that X-linked expression is somewhat elevated in reproductive tracts of adult males as compared to those of 4^th^ nymphal stage males (Figure 3A). This is most likely due to the larger portion of somatic tissues in adult male reproductive tracts, most notably accessory glands, which are very small in all nymphal stages but well developed in adults (Jelisaveta Djordjevic, personal observation). Such a pattern is known to also occur in *Caenorhabditis elegans*, where the X is completely silenced in germ cells and changes in X-linked expression during development are caused by changes in the ratio of somatic to germ cells [52, 72, 73]. More generally, the observation of reduced X-expression in male gonads without investigation of associated mechanism is typically interpreted as lack of dosage compensation in this tissue [11, 12, 52, 74]. However, as we observe in *Timema*, dosage compensation in males may well be ubiquitous across tissues and developmental stages in many species, and reduced X-expression patterns would solely be driven by MSCI.

## Conclusion

Dosage compensation by upregulation of the single X in males consistently occurs in somatic tissues across development in *Timema poppense*. By contrast, male X-expression in the reproductive tract is dynamic. It starts with complete dosage compensation in the first nymphal stage, when spermatogonia have not yet entered meiosis I, and is followed by X inactivation in later nymphal stages when most gonadal cells are undergoing meiotic divisions. We suggest that the reduced X-linked expression in testes is solely driven by X chromosome inactivation during meiosis, and does not stem from a lack of dosage compensation.

## Acknowledgements

We thank Corinne Peter and the Lausanne GTF platform for help with ChIP-seq, Bart Zijlstra, Armand Yazdani, Susana Freitas, and Chloé Larose for help in the field, and current and previous members of the Schwander lab for discussions. Funding: We would like to acknowledge funding from the European Research Council Consolidator Grant (No Sex No Conflict to T.S.) and Swiss FNS grant 31003A_182495 (T.S.) and support from the Genoscope, the Commissariat à l’Énergie Atomique et aux Énergies Alternatives (CEA) and France Génomique (ANR-10-INBS-09–08). Author contributions: J.D. and T.S. designed the study. J.D., Z.D., M.L. and W.T, C.C. and K.L. performed molecular work. W.T., Z.D., J-M.A., B.I. and B.N. developed methods J. D., D.J.P., and P.T.V. analyzed the data with input from T.S.. J.D. and T.S. wrote the paper with input from all authors. Competing interests: The authors declare that they have no competing interests. Data and code availability: Genome assembly: PRJNA1126215; annotation (https://github.com/JelisavetaDjordjevic/Dosage_compensation/tree/main/Annotation); raw sequence reads have been deposited in NCBI’s sequence read archive under the following bioprojects: PRJEB76439 (reference genome), PRJNA1128554 (RNAseq reads). Data were processed to generate plots and statistics using R v3.4.4.

## Bibliography

1. Stingele S, Stoehr G, Peplowska K, Cox J, Mann M, Storchova Z. Global analysis of genome, transcriptome and proteome reveals the response to aneuploidy in human cells. Molecular Systems Biology. 2012;8(1):608. doi: 10.1038/msb.2012.40.

2. Ohno S. Sex chromosomes and sex-linked genes. Verlag: Springer; 1967.

3. Straub T, Becker PB. Dosage compensation: the beginning and end of generalization. Nature Reviews Genetics. 2007;8(1):47–57.

4. Mank JE. Sex chromosome dosage compensation: definitely not for everyone. Trends in genetics. 2013;29(12):677–83.

5. Gupta V, Parisi M, Sturgill D, Nuttall R, Doctolero M, Dudko OK, et al. Global analysis of X-chromosome dosage compensation. Journal of Biology. 2006;5(1):3. doi: 10.1186/jbiol30.

6. Nguyen DK, Disteche CM. Dosage compensation of the active X chromosome in mammals. Nature Genetics. 2006;38(1):47–53. doi: 10.1038/ng1705.

7. Lyon MF. Gene Action in the X-chromosome of the Mouse (Mus musculus L.). Nature. 1961;190(4773):372-3. doi: 10.1038/190372a0.

8. Meyer BJ, Casson LP. Caenorhabditis elegans compensates for the difference in X chromosome dosage between the sexes by regulating transcript levels. Cell. 1986;47(6):871–81.

9. Meyer B, McDonel P, Csankovszki G, Ralston E, editors. Sex and X-Chromosome-wide Repression in Caenorhabditiselegans. Cold Spring Harbor symposia on quantitative biology; 2004: Cold Spring Harbor Laboratory Press.

10. Lucchesi JC. Dosage compensation in flies and worms: the ups and downs of X-chromosome regulation. Current Opinion in Genetics & Development. 1998;8(2):179–84. doi: 10.1016/S0959-437X(98)80139-1.

11. Vicoso B, Bachtrog D. Numerous Transitions of Sex Chromosomes in Diptera. PLOS Biology. 2015;13(4):e1002078. doi: 10.1371/journal.pbio.1002078.

12. Hu Q-L, Ye Y-X, Zhuo J-C, Huang H-J, Li J-M, Zhang C-X. Chromosome-level Assembly, Dosage Compensation and Sex-biased Gene Expression in the Small Brown Planthopper, Laodelphax striatellus. Genome Biology and Evolution. 2022;14(11). doi: 10.1093/gbe/evac160.

13. Parker DJ, Jaron KS, Dumas Z, Robinson-Rechavi M, Schwander T. X chromosomes show relaxed selection and complete somatic dosage compensation across Timema stick insect species. Journal of Evolutionary Biology. 2022;35(12):1734–50. doi: 10.1111/jeb.14075.

14. Blackmon H, Ross L, Bachtrog D. Sex determination, sex chromosomes, and karyotype evolution in insects. Journal of Heredity. 2017;108(1):78–93.

15. Toups MA, Vicoso B. The X chromosome of insects predates the origin of Class Insecta. bioRxiv. 2023:2023.04.19.537501. doi: 10.1101/2023.04.19.537501.

16. Pease JB, Hahn MW. Sex chromosomes evolved from independent ancestral linkage groups in winged insects. Molecular biology and evolution. 2012;29(6):1645–53.

17. Hamada FN, Park PJ, Gordadze PR, Kuroda MI. Global regulation of X chromosomal genes by the MSL complex in Drosophila melanogaster. Genes & development. 2005;19(19):2289–94.

18. Straub T, Gilfillan GD, Maier VK, Becker PB. The Drosophila MSL complex activates the transcription of target genes. Genes & development. 2005;19(19):2284–8.

19. Kalita AI, Marois E, Kozielska M, Weissing FJ, Jaouen E, Möckel MM, et al. The sex-specific factor SOA controls dosage compensation in Anopheles mosquitoes. Nature. 2023;623(7985):175–82.

20. Krzywinska E, Ribeca P, Ferretti L, Hammond A, Krzywinski J. A novel factor modulating X chromosome dosage compensation in Anopheles. Current Biology. 2023;33(21):4697–703. e4.

21. Rayner JG, Hitchcock TJ, Bailey NW. Variable dosage compensation is associated with female consequences of an X-linked, male-beneficial mutation. Proceedings of the Royal Society B. 2021;288(1947):20210355.

22. Turner JMA. Meiotic sex chromosome inactivation. Development. 2007;134(10):1823–31. doi: 10.1242/dev.000018.

23. Namekawa SH, Lee JT. XY and ZW: is meiotic sex chromosome inactivation the rule in evolution? PLoS genetics. 2009;5(5):e1000493.

24. Kramer M, Kranz A-L, Su A, Winterkorn LH, Albritton SE, Ercan S. Developmental dynamics of X-chromosome dosage compensation by the DCC and H4K20me1 in C. elegans. PLoS genetics. 2015;11(12):e1005698.

25. Schwander T, Crespi BJ. Multiple direct transitions from sexual reproduction to apomictic parthenogenesis in Timema stick insects. Evolution. 2009;63(1):84–103.

26. Kolmogorov M, Yuan J, Lin Y, Pevzner PA. Assembly of long, error-prone reads using repeat graphs. Nature biotechnology. 2019;37(5):540–6.

27. Li H. Minimap2: pairwise alignment for nucleotide sequences. Bioinformatics. 2018;34(18):3094–100.

28. Vaser R, Sović I, Nagarajan N, Šikić M. Fast and accurate de novo genome assembly from long uncorrected reads. Genome research. 2017;27(5):737–46.

29. Li H. Aligning sequence reads, clone sequences and assembly contigs with BWA-MEM. arXiv preprint arXiv:13033997. 2013.

30. Walker BJ, Abeel T, Shea T, Priest M, Abouelliel A, Sakthikumar S, et al. Pilon: an integrated tool for comprehensive microbial variant detection and genome assembly improvement. PloS one. 2014;9(11):e112963.

31. Laetsch DR, Blaxter ML. BlobTools: Interrogation of genome assemblies. F1000Research. 2017;6:1287.

32. Roach MJ, Schmidt SA, Borneman AR. Purge Haplotigs: allelic contig reassignment for third-gen diploid genome assemblies. BMC bioinformatics. 2018;19:1–10.

33. Durand NC, Shamim MS, Machol I, Rao SS, Huntley MH, Lander ES, et al. Juicer provides a one-click system for analyzing loop-resolution Hi-C experiments. Cell systems. 2016;3(1):95–8.

34. Dudchenko O, Batra SS, Omer AD, Nyquist SK, Hoeger M, Durand NC, et al. De novo assembly of the Aedes aegypti genome using Hi-C yields chromosome-length scaffolds. Science. 2017;356(6333):92–5.

35. Seppey M, Manni M, Zdobnov EM. BUSCO: assessing genome assembly and annotation completeness. Gene prediction: methods and protocols. 2019:227–45.

36. Jaron KS, Parker DJ, Anselmetti Y, Tran Van P, Bast J, Dumas Z, et al. Convergent consequences of parthenogenesis on stick insect genomes. Science Advances. 2022;8(8):eabg3842. doi: doi:10.1126/sciadv.abg3842.

37. Brůna T, Hoff KJ, Lomsadze A, Stanke M, Borodovsky M. BRAKER2: automatic eukaryotic genome annotation with GeneMark-EP+ and AUGUSTUS supported by a protein database. NAR genomics and bioinformatics. 2021;3(1):lqaa108.

38. Kriventseva EV, Kuznetsov D, Tegenfeldt F, Manni M, Dias R, Simão FA, et al. OrthoDB v10: sampling the diversity of animal, plant, fungal, protist, bacterial and viral genomes for evolutionary and functional annotations of orthologs. Nucleic acids research. 2019;47(D1):D807–D11.

39. Bolger AM, Lohse M, Usadel B. Trimmomatic: a flexible trimmer for Illumina sequence data. Bioinformatics. 2014;30(15):2114–20.

40. Dobin A, Davis CA, Schlesinger F, Drenkow J, Zaleski C, Jha S, et al. STAR: ultrafast universal RNA-seq aligner. Bioinformatics. 2013;29(1):15–21. Epub 2012/10/30. doi: 10.1093/bioinformatics/bts635. PubMed PMID: 23104886; PubMed Central PMCID: PMCPMC3530905.

41. Stanke M, Keller O, Gunduz I, Hayes A, Waack S, Morgenstern B. AUGUSTUS: ab initio prediction of alternative transcripts. Nucleic acids research. 2006;34(suppl_2):W435–W9.

42. Brůna T, Lomsadze A, Borodovsky M. GeneMark-EP+: eukaryotic gene prediction with self-training in the space of genes and proteins. NAR genomics and bioinformatics. 2020;2(2):lqaa026.

43. Keilwagen J, Hartung F, Grau J. GeMoMa: homology-based gene prediction utilizing intron position conservation and RNA-seq data. Gene prediction: Methods and protocols. 2019:161–77.

44. Gabriel L, Hoff KJ, Brůna T, Borodovsky M, Stanke M. TSEBRA: transcript selector for BRAKER. Bmc Bioinformatics. 2021;22:1–12.

45. Nawrocki EP, Eddy SR. Infernal 1.1: 100-fold faster RNA homology searches. Bioinformatics. 2013;29(22):2933–5.

46. Djordjevic J, Dumas Z, Robinson-Rechavi M, Schwander T, Parker DJ. Dynamics of sex-biased gene expression during development in the stick insect Timema californicum. Heredity. 2022:1–10.

47. Anders S, Pyl PT, Huber W. HTSeq—a Python framework to work with high-throughput sequencing data. Bioinformatics. 2014;31(2):166–9. doi: 10.1093/bioinformatics/btu638.

48. Lawrence M, Huber W, Pagès H, Aboyoun P, Carlson M, Gentleman R, et al. Software for Computing and Annotating Genomic Ranges. PLOS Computational Biology. 2013;9(8):e1003118. doi: 10.1371/journal.pcbi.1003118.

49. Robinson MD, McCarthy DJ, Smyth GK. edgeR: a Bioconductor package for differential expression analysis of digital gene expression data. Bioinformatics. 2009;26(1):139–40. doi: 10.1093/bioinformatics/btp616.

50. Yanai I, Benjamin H, Shmoish M, Chalifa-Caspi V, Shklar M, Ophir R, et al. Genome-wide midrange transcription profiles reveal expression level relationships in human tissue specification. Bioinformatics. 2004;21(5):650–9. doi: 10.1093/bioinformatics/bti042.

51. Wickham H. ggplot2: Elegant Graphics for Data Analysis: Springer-Verlag New York; 2016.

52. Gu L, Walters JR. Evolution of Sex Chromosome Dosage Compensation in Animals: A Beautiful Theory, Undermined by Facts and Bedeviled by Details. Genome Biology and Evolution. 2017;9(9):2461–76. doi: 10.1093/gbe/evx154.

53. Stuart OP, Cleave R, Magrath MJL, Mikheyev AS. Genome of the Lord Howe Island Stick Insect Reveals a Highly Conserved Phasmid X Chromosome. Genome Biology and Evolution. 2023;15(6). doi: 10.1093/gbe/evad104.

54. Vicoso B, Charlesworth B. Effective population size and the faster-X effect: an extended model. Evolution. 2009;63(9):2413–26.

55. Hurst LD, Ghanbarian AT, Forrest ARR, consortium F, Huminiecki L. The Constrained Maximal Expression Level Owing to Haploidy Shapes Gene Content on the Mammalian X Chromosome. PLOS Biology. 2015;13(12):e1002315. doi: 10.1371/journal.pbio.1002315.

56. Argyridou E, Parsch J. Regulation of the X Chromosome in the Germline and Soma of Drosophila melanogaster Males. Genes. 2018;9(5):242. PubMed PMID: doi:10.3390/genes9050242.

57. Pessia E, Engelstädter J, Marais GAB. The evolution of X chromosome inactivation in mammals: the demise of Ohno’s hypothesis? Cellular and Molecular Life Sciences. 2014;71(8):1383–94. doi: 10.1007/s00018-013-1499-6.

58. Bachtrog D. Y-chromosome evolution: emerging insights into processes of Y-chromosome degeneration. Nature Reviews Genetics. 2013;14(2):113–24. doi: 10.1038/nrg3366.

59. Turner JM. Meiotic silencing in mammals. Annual review of genetics. 2015;49:395–412.

60. Viera A, Parra MT, Arévalo S, García de la Vega C, Santos JL, Page J. X Chromosome Inactivation during Grasshopper Spermatogenesis. Genes (Basel). 2021;12(12). Epub 2021/12/25. doi: 10.3390/genes12121844. PubMed PMID: 34946793; PubMed Central PMCID: PMCPMC8700825.

61. Mahadevaraju S, Fear JM, Akeju M, Galletta BJ, Pinheiro MM, Avelino CC, et al. Dynamic sex chromosome expression in Drosophila male germ cells. Nature communications. 2021;12(1):892.

62. Witt E, Shao Z, Hu C, Krause HM, Zhao L. Single-cell RNA-sequencing reveals pre-meiotic X-chromosome dosage compensation in Drosophila testis. PLoS genetics. 2021;17(8):e1009728.

63. Raz AA, Vida GS, Stern SR, Mahadevaraju S, Fingerhut JM, Viveiros JM, et al. Emergent dynamics of adult stem cell lineages from single nucleus and single cell RNA-Seq of Drosophila testes. Elife. 2023;12:e82201.

64. Anderson J, Henikoff S, Ahmad K. Chromosome-specific maturation of the epigenome in the Drosophila male germline. eLife Sciences Publications, Ltd; 2023.

65. Schoenmakers S, Wassenaar E, Hoogerbrugge JW, Laven JSE, Grootegoed JA, Baarends WM. Female Meiotic Sex Chromosome Inactivation in Chicken. PLOS Genetics. 2009;5(5):e1000466. doi: 10.1371/journal.pgen.1000466.

66. McKee BD, Handel MA. Sex chromosomes, recombination, and chromatin conformation. Chromosoma. 1993;102(2):71–80.

67. Kelly WG, Aramayo R. Meiotic silencing and the epigenetics of sex. Chromosome research. 2007;15:633–51.

68. Meiklejohn CD, Tao Y. Genetic conflict and sex chromosome evolution. Trends in ecology & evolution. 2010;25(4):215–23.

69. Wu C-I, Xu EY. Sexual antagonism and X inactivation–the SAXI hypothesis. TRENDS in Genetics. 2003;19(5):243–7.

70. Vicoso B, Charlesworth B. Evolution on the X chromosome: unusual patterns and processes. Nature Reviews Genetics. 2006;7(8):645–53.

71. McKee BD, Habera L, Vrana JA. Evidence that intergenic spacer repeats of Drosophila melanogaster rRNA genes function as X-Y pairing sites in male meiosis, and a general model for achiasmatic pairing. Genetics. 1992;132(2):529–44. doi: 10.1093/genetics/132.2.529.

72. Kelly WG, Schaner CE, Dernburg AF, Lee M-H, Kim SK, Villeneuve AM, et al. X-chromosome silencing in the germline of C. elegans. Development. 2002;129(2):479–92. doi: 10.1242/dev.129.2.479.

73. Deng X, Hiatt JB, Nguyen DK, Ercan S, Sturgill D, Hillier LW, et al. Evidence for compensatory upregulation of expressed X-linked genes in mammals, Caenorhabditis elegans and Drosophila melanogaster. Nature Genetics. 2011;43(12):1179–85. doi: 10.1038/ng.948.

74. Parker DJ, Jaron KS, Dumas Z, Robinson-Rechavi M, Schwander T. X chromosomes show relaxed selection and complete somatic dosage compensation across Timema stick insect species. Journal of Evolutionary Biology. 2022;35(12):1734–50.

