## Supplemental material for "Dynamics of X chromosome hyper-expression and inactivation in male tissues during stick insect development"

**Supplemental Tables**

**Supplemental Table S1.** Number of replicates per sex, for every developmental stage and tissue

|  | **Adult** | | **Nymphal 1** | | **Nymphal 2** | | **Nymphal 3** | | **Nymphal 4** | |
| --- | --- | --- | --- | --- | --- | --- | --- | --- | --- | --- |
| **Tissue/Sex** | F | M | F | M | F | M | F | M | F | M |
| **Antennae** | 4 | 4 | 3 | 2 | 3 | 3 | 2 | 3 | 3 | 3 |
| **Brain** | 4 | 4 | 3 | 3 | 3 | 3 | 2 | 3 | 3 | 3 |
| **Gonads** | 4 | 3 | 4 | 4 | 0 | 0 | 0 | 0 | 3 | 3 |
| **Guts** | 4 | 4 | 3 | 3 | 3 | 3 | 2 | 3 | 2 | 3 |
| **Legs** | 0 | 0 | 3 | 3 | 3 | 3 | 2 | 3 | 3 | 3 |

**Supplemental Table S2**. Results of Wilcoxon tests where the ratio of male to female average RPKM values were compared between autosomes and the X chromosome within every developmental stage and tissue. The column “stage” indicates the developmental stage; N1-N4 (1^st^ to 4^th^ nymphal stage), A- adult, the column “tissue” the different tissues, the column; “p.adjusted”- fdr correction of p-values for multiple testing using the Benjamini & Hochberg method (Benjamini and Hochberg 1995)*.*

| **Stage** | **Group1** | **Group2** | **p.adjusted** | **Tissue** |
| --- | --- | --- | --- | --- |
| A | Autosome | X chromosome | 4.80E-49 | brain |
| N1 | Autosome | X chromosome | 2.10E-26 | brain |
| N3 | Autosome | X chromosome | 0.73 | brain |
| N2 | Autosome | X chromosome | 0.021 | brain |
| N4 | Autosome | X chromosome | 2.50E-18 | brain |
| A | Autosome | X chromosome | 3.90E-19 | antenna |
| N1 | Autosome | X chromosome | 1.30E-14 | antenna |
| N3 | Autosome | X chromosome | 1.80E-26 | antenna |
| N2 | Autosome | X chromosome | 0.0027 | antenna |
| N4 | Autosome | X chromosome | 1.30E-18 | antenna |
| A | Autosome | X chromosome | 0.055 | gut |
| N1 | Autosome | X chromosome | 3.80E-14 | gut |
| N3 | Autosome | X chromosome | 0.95 | gut |
| N2 | Autosome | X chromosome | 0.065 | gut |
| N4 | Autosome | X chromosome | 0.067 | gut |
| N1 | Autosome | X chromosome | 9.80E-12 | leg |
| N3 | Autosome | X chromosome | 0.00085 | leg |
| N2 | Autosome | X chromosome | 0.0077 | leg |
| N4 | Autosome | X chromosome | 1.20E-13 | leg |
| A | Autosome | X chromosome | 2.10E-176 | gonad |
| N1 | Autosome | X chromosome | 1.60E-06 | gonad |
| N4 | Autosome | X chromosome | 1.30E-247 | gonad |

**Supplemental Table S3.** Results of Wilcoxon tests where average RPKM values were compared for autosomal and X-linked genes for each sex and within each developmental stage and tissue (indicated in the "Tissue" column). The "Stage" column denotes the developmental stage, with N1-N4 representing the 1st to 4th nymphal stages and A indicating the adult stage. **Columns: Group1 and Group2:** Specify the groups being compared, with colored cells highlighting the comparisons of interest. **"p.adjusted":** Represents the false discovery rate (FDR) corrected p-values, utilizing the Benjamini & Hochberg method for multiple testing.

| **Stage** | **Group1** | **Group2** | **p.adjusted** | **Tissue** |
| --- | --- | --- | --- | --- |
| A | Autosomes_F | Autosomes_M | 0.76 | brain |
| A | Autosomes_F | X_chromosome_F | 0.23 | brain |
| A | Autosomes_F | X_chromosome_M | 0.32 | brain |
| A | Autosomes_M | X_chromosome_F | 0.17 | brain |
| A | Autosomes_M | X_chromosome_M | 0.41 | brain |
| A | X_chromosome_F | X_chromosome_M | 0.11 | brain |
| N1 | Autosomes_F | Autosomes_M | 0.99 | brain |
| N1 | Autosomes_F | X_chromosome_F | 0.99 | brain |
| N1 | Autosomes_F | X_chromosome_M | 0.097 | brain |
| N1 | Autosomes_M | X_chromosome_F | 0.99 | brain |
| N1 | Autosomes_M | X_chromosome_M | 0.097 | brain |
| N1 | X_chromosome_F | X_chromosome_M | 0.17 | brain |
| N3 | Autosomes_F | Autosomes_M | 9.20E-32 | brain |
| N3 | Autosomes_F | X_chromosome_F | 0.11 | brain |
| N3 | Autosomes_F | X_chromosome_M | 0.094 | brain |
| N3 | Autosomes_M | X_chromosome_F | 1.50E-10 | brain |
| N3 | Autosomes_M | X_chromosome_M | 0.094 | brain |
| N3 | X_chromosome_F | X_chromosome_M | 0.0062 | brain |
| N2 | Autosomes_F | Autosomes_M | 0.99 | brain |
| N2 | Autosomes_F | X_chromosome_F | 0.76 | brain |
| N2 | Autosomes_F | X_chromosome_M | 0.11 | brain |
| N2 | Autosomes_M | X_chromosome_F | 0.76 | brain |
| N2 | Autosomes_M | X_chromosome_M | 0.11 | brain |
| N2 | X_chromosome_F | X_chromosome_M | 0.41 | brain |
| N4 | Autosomes_F | Autosomes_M | 0.99 | brain |
| N4 | Autosomes_F | X_chromosome_F | 0.71 | brain |
| N4 | Autosomes_F | X_chromosome_M | 0.15 | brain |
| N4 | Autosomes_M | X_chromosome_F | 0.7 | brain |
| N4 | Autosomes_M | X_chromosome_M | 0.15 | brain |
| N4 | X_chromosome_F | X_chromosome_M | 0.56 | brain |
| A | Autosomes_F | Autosomes_M | 0.59 | antenna |
| A | Autosomes_F | X_chromosome_F | 0.049* | antenna |
| A | Autosomes_F | X_chromosome_M | 0.59 | antenna |
| A | Autosomes_M | X_chromosome_F | 0.038 | antenna |
| A | Autosomes_M | X_chromosome_M | 0.75 | antenna |
| A | X_chromosome_F | X_chromosome_M | 0.058 | antenna |
| N1 | Autosomes_F | Autosomes_M | 0.69 | antenna |
| N1 | Autosomes_F | X_chromosome_F | 0.91 | antenna |
| N1 | Autosomes_F | X_chromosome_M | 0.066 | antenna |
| N1 | Autosomes_M | X_chromosome_F | 0.95 | antenna |
| N1 | Autosomes_M | X_chromosome_M | 0.049* | antenna |
| N1 | X_chromosome_F | X_chromosome_M | 0.14 | antenna |
| N3 | Autosomes_F | Autosomes_M | 0.93 | antenna |
| N3 | Autosomes_F | X_chromosome_F | 0.59 | antenna |
| N3 | Autosomes_F | X_chromosome_M | 0.00038 | antenna |
| N3 | Autosomes_M | X_chromosome_F | 0.59 | antenna |
| N3 | Autosomes_M | X_chromosome_M | 0.00038* | antenna |
| N3 | X_chromosome_F | X_chromosome_M | 0.039* | antenna |
| N2 | Autosomes_F | Autosomes_M | 0.76 | antenna |
| N2 | Autosomes_F | X_chromosome_F | 0.088 | antenna |
| N2 | Autosomes_F | X_chromosome_M | 0.0013 | antenna |
| N2 | Autosomes_M | X_chromosome_F | 0.12 | antenna |
| N2 | Autosomes_M | X_chromosome_M | 0.0021* | antenna |
| N2 | X_chromosome_F | X_chromosome_M | 0.33 | antenna |
| N4 | Autosomes_F | Autosomes_M | 0.52 | antenna |
| N4 | Autosomes_F | X_chromosome_F | 0.063 | antenna |
| N4 | Autosomes_F | X_chromosome_M | 0.001 | antenna |
| N4 | Autosomes_M | X_chromosome_F | 0.038 | antenna |
| N4 | Autosomes_M | X_chromosome_M | 0.00038* | antenna |
| N4 | X_chromosome_F | X_chromosome_M | 0.36 | antenna |
| A | Autosomes_F | Autosomes_M | 0.21 | gut |
| A | Autosomes_F | X_chromosome_F | 0.00048* | gut |
| A | Autosomes_F | X_chromosome_M | 3.00E-04 | gut |
| A | Autosomes_M | X_chromosome_F | 4.10E-05 | gut |
| A | Autosomes_M | X_chromosome_M | 2.50E-05* | gut |
| A | X_chromosome_F | X_chromosome_M | 0.97 | gut |
| N1 | Autosomes_F | Autosomes_M | 0.13 | gut |
| N1 | Autosomes_F | X_chromosome_F | 0.0017* | gut |
| N1 | Autosomes_F | X_chromosome_M | 1.40E-05 | gut |
| N1 | Autosomes_M | X_chromosome_F | 0.00013 | gut |
| N1 | Autosomes_M | X_chromosome_M | 5.00E-07* | gut |
| N1 | X_chromosome_F | X_chromosome_M | 0.4 | gut |
| N3 | Autosomes_F | Autosomes_M | 6.10E-08 | gut |
| N3 | Autosomes_F | X_chromosome_F | 4.70E-06* | gut |
| N3 | Autosomes_F | X_chromosome_M | 0.0046 | gut |
| N3 | Autosomes_M | X_chromosome_F | 5.30E-12 | gut |
| N3 | Autosomes_M | X_chromosome_M | 2.50E-07* | gut |
| N3 | X_chromosome_F | X_chromosome_M | 0.2 | gut |
| N2 | Autosomes_F | Autosomes_M | 0.46 | gut |
| N2 | Autosomes_F | X_chromosome_F | 4.30E-05* | gut |
| N2 | Autosomes_F | X_chromosome_M | 5.00E-07 | gut |
| N2 | Autosomes_M | X_chromosome_F | 1.30E-05 | gut |
| N2 | Autosomes_M | X_chromosome_M | 1.20E-07* | gut |
| N2 | X_chromosome_F | X_chromosome_M | 0.47 | gut |
| N4 | Autosomes_F | Autosomes_M | 0.3 | gut |
| N4 | Autosomes_F | X_chromosome_F | 0.99 | gut |
| N4 | Autosomes_F | X_chromosome_M | 0.75 | gut |
| N4 | Autosomes_M | X_chromosome_F | 0.75 | gut |
| N4 | Autosomes_M | X_chromosome_M | 0.47 | gut |
| N4 | X_chromosome_F | X_chromosome_M | 0.76 | gut |
| N1 | Autosomes_F | Autosomes_M | 0.092 | leg |
| N1 | Autosomes_F | X_chromosome_F | 0.29 | leg |
| N1 | Autosomes_F | X_chromosome_M | 0.018 | leg |
| N1 | Autosomes_M | X_chromosome_F | 0.067 | leg |
| N1 | Autosomes_M | X_chromosome_M | 0.0015* | leg |
| N1 | X_chromosome_F | X_chromosome_M | 0.35 | leg |
| N3 | Autosomes_F | Autosomes_M | 0.53 | leg |
| N3 | Autosomes_F | X_chromosome_F | 0.4 | leg |
| N3 | Autosomes_F | X_chromosome_M | 0.19 | leg |
| N3 | Autosomes_M | X_chromosome_F | 0.34 | leg |
| N3 | Autosomes_M | X_chromosome_M | 0.11 | leg |
| N3 | X_chromosome_F | X_chromosome_M | 0.59 | leg |
| N2 | Autosomes_F | Autosomes_M | 0.46 | leg |
| N2 | Autosomes_F | X_chromosome_F | 0.45 | leg |
| N2 | Autosomes_F | X_chromosome_M | 0.059 | leg |
| N2 | Autosomes_M | X_chromosome_F | 0.59 | leg |
| N2 | Autosomes_M | X_chromosome_M | 0.096 | leg |
| N2 | X_chromosome_F | X_chromosome_M | 0.35 | leg |
| N4 | Autosomes_F | Autosomes_M | 0.59 | leg |
| N4 | Autosomes_F | X_chromosome_F | 0.31 | leg |
| N4 | Autosomes_F | X_chromosome_M | 0.018 | leg |
| N4 | Autosomes_M | X_chromosome_F | 0.36 | leg |
| N4 | Autosomes_M | X_chromosome_M | 0.034* | leg |
| N4 | X_chromosome_F | X_chromosome_M | 0.35 | leg |
| A | Autosomes_F | Autosomes_M | 2.70E-87 | gonad |
| A | Autosomes_F | X_chromosome_F | 7.70E-07* | gonad |
| A | Autosomes_F | X_chromosome_M | 7.20E-25 | gonad |
| A | Autosomes_M | X_chromosome_F | 0.014 | gonad |
| A | Autosomes_M | X_chromosome_M | 9.70E-80* | gonad |
| A | X_chromosome_F | X_chromosome_M | 1.60E-36* | gonad |
| N1 | Autosomes_F | Autosomes_M | 0.88 | gonad |
| N1 | Autosomes_F | X_chromosome_F | 0.0017* | gonad |
| N1 | Autosomes_F | X_chromosome_M | 4.70E-08 | gonad |
| N1 | Autosomes_M | X_chromosome_F | 0.0022 | gonad |
| N1 | Autosomes_M | X_chromosome_M | 9.50E-08* | gonad |
| N1 | X_chromosome_F | X_chromosome_M | 0.075 | gonad |
| N4 | Autosomes_F | Autosomes_M | 3.50E-55 | gonad |
| N4 | Autosomes_F | X_chromosome_F | 0.033* | gonad |
| N4 | Autosomes_F | X_chromosome_M | 3.50E-64 | gonad |
| N4 | Autosomes_M | X_chromosome_F | 0.00078 | gonad |
| N4 | Autosomes_M | X_chromosome_M | 1.20E-127* | gonad |
| N4 | X_chromosome_F | X_chromosome_M | 2.50E-64 | gonad |

**Supplemental Table S4.** Sex-biased gene counts during reproductive tract development at N1 (1st nymphal), N4 (4th nymphal), and A (adult) stages. It includes MB (male-biased), FB (female-biased), SB (total sex-biased genes), and T (total expressed genes) categories, with percentages in brackets. Differential gene expression between the sexes was assessed using a generalized linear model with a quasi-likelihood F-test (Chen, et al. 2016) in edgeR v.3.42.4 (Robinson, et al. 2009; McCarthy, et al. 2012) in each of the three developmental stages as described in (Djordjevic, et al. 2022).

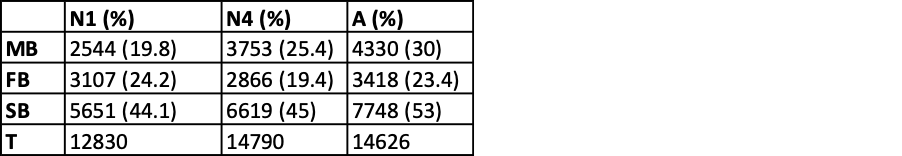

**Supplemental Table S5.** Wilcoxon tests comparing tissue specificity [Tau], calculated across three somatic tissues, between scaffolds within each sex **(“variable”**; Females, Males). **Columns: Group1 and Group2:** Specify the scaffolds being compared, with colored cells highlighting the comparisons of interest, **“p.adjusted”:** Represents the false discovery rate (FDR) corrected p-values, utilizing the Benjamini & Hochberg method for multiple testing.

| **Variable** | **Group1** | **Group2** | **p.adjusted** |
| --- | --- | --- | --- |
| Female | Tps_LRv5b_scf1 | Tps_LRv5b_scf2 | 0.016 |
| Female | Tps_LRv5b_scf1 | Tps_LRv5b_scf3 | 8.10E-16 |
| Female | Tps_LRv5b_scf1 | Tps_LRv5b_scf4 | 0.0031 |
| Female | Tps_LRv5b_scf1 | Tps_LRv5b_scf5 | 4.00E-04 |
| Female | Tps_LRv5b_scf1 | Tps_LRv5b_scf6 | 0.29 |
| Female | Tps_LRv5b_scf1 | Tps_LRv5b_scf7 | 0.16 |
| Female | Tps_LRv5b_scf1 | Tps_LRv5b_scf8 | 0.0066 |
| Female | Tps_LRv5b_scf1 | Tps_LRv5b_scf9 | 0.25 |
| Female | Tps_LRv5b_scf1 | Tps_LRv5b_scf10 | 0.0026 |
| Female | Tps_LRv5b_scf1 | Tps_LRv5b_scf11 | 5.60E-10 |
| Female | Tps_LRv5b_scf1 | Tps_LRv5b_scf12 | 0.17 |
| Female | Tps_LRv5b_scf2 | Tps_LRv5b_scf3 | 2.30E-07 |
| Female | Tps_LRv5b_scf2 | Tps_LRv5b_scf4 | 0.5 |
| Female | Tps_LRv5b_scf2 | Tps_LRv5b_scf5 | 0.22 |
| Female | Tps_LRv5b_scf2 | Tps_LRv5b_scf6 | 0.46 |
| Female | Tps_LRv5b_scf2 | Tps_LRv5b_scf7 | 0.65 |
| Female | Tps_LRv5b_scf2 | Tps_LRv5b_scf8 | 0.46 |
| Female | Tps_LRv5b_scf2 | Tps_LRv5b_scf9 | 0.005 |
| Female | Tps_LRv5b_scf2 | Tps_LRv5b_scf10 | 8.00E-06 |
| Female | Tps_LRv5b_scf2 | Tps_LRv5b_scf11 | 0.00015 |
| Female | Tps_LRv5b_scf2 | Tps_LRv5b_scf12 | 0.92 |
| Female | Tps_LRv5b_scf3 | Tps_LRv5b_scf4 | 6.30E-05 |
| Female | Tps_LRv5b_scf3 | Tps_LRv5b_scf5 | 0.0013 |
| Female | Tps_LRv5b_scf3 | Tps_LRv5b_scf6 | 2.20E-07 |
| Female | Tps_LRv5b_scf3 | Tps_LRv5b_scf7 | 2.30E-06 |
| Female | Tps_LRv5b_scf3 | Tps_LRv5b_scf8 | 0.00045 |
| Female | Tps_LRv5b_scf3 | Tps_LRv5b_scf9 | 1.20E-10 |
| Female | Tps_LRv5b_scf3 | Tps_LRv5b_scf10 | 2.20E-17 |
| Female | Tps_LRv5b_scf3 | Tps_LRv5b_scf11 | 0.5 |
| Female | Tps_LRv5b_scf3 | Tps_LRv5b_scf12 | 0.00012 |
| Female | Tps_LRv5b_scf4 | Tps_LRv5b_scf5 | 0.59 |
| Female | Tps_LRv5b_scf4 | Tps_LRv5b_scf6 | 0.21 |
| Female | Tps_LRv5b_scf4 | Tps_LRv5b_scf7 | 0.33 |
| Female | Tps_LRv5b_scf4 | Tps_LRv5b_scf8 | 0.9 |
| Female | Tps_LRv5b_scf4 | Tps_LRv5b_scf9 | 0.0021 |
| Female | Tps_LRv5b_scf4 | Tps_LRv5b_scf10 | 2.30E-06 |
| Female | Tps_LRv5b_scf4 | Tps_LRv5b_scf11 | 0.0035 |
| Female | Tps_LRv5b_scf4 | Tps_LRv5b_scf12 | 0.55 |
| Female | Tps_LRv5b_scf5 | Tps_LRv5b_scf6 | 0.083 |
| Female | Tps_LRv5b_scf5 | Tps_LRv5b_scf7 | 0.15 |
| Female | Tps_LRv5b_scf5 | Tps_LRv5b_scf8 | 0.74 |
| Female | Tps_LRv5b_scf5 | Tps_LRv5b_scf9 | 0.00056 |
| Female | Tps_LRv5b_scf5 | Tps_LRv5b_scf10 | 3.80E-07 |
| Female | Tps_LRv5b_scf5 | Tps_LRv5b_scf11 | 0.024 |
| Female | Tps_LRv5b_scf5 | Tps_LRv5b_scf12 | 0.3 |
| Female | Tps_LRv5b_scf6 | Tps_LRv5b_scf7 | 0.85 |
| Female | Tps_LRv5b_scf6 | Tps_LRv5b_scf8 | 0.2 |
| Female | Tps_LRv5b_scf6 | Tps_LRv5b_scf9 | 0.064 |
| Female | Tps_LRv5b_scf6 | Tps_LRv5b_scf10 | 0.001 |
| Female | Tps_LRv5b_scf6 | Tps_LRv5b_scf11 | 6.40E-05 |
| Female | Tps_LRv5b_scf6 | Tps_LRv5b_scf12 | 0.66 |
| Female | Tps_LRv5b_scf7 | Tps_LRv5b_scf8 | 0.3 |
| Female | Tps_LRv5b_scf7 | Tps_LRv5b_scf9 | 0.052 |
| Female | Tps_LRv5b_scf7 | Tps_LRv5b_scf10 | 0.00062 |
| Female | Tps_LRv5b_scf7 | Tps_LRv5b_scf11 | 0.00026 |
| Female | Tps_LRv5b_scf7 | Tps_LRv5b_scf12 | 0.89 |
| Female | Tps_LRv5b_scf8 | Tps_LRv5b_scf9 | 0.0025 |
| Female | Tps_LRv5b_scf8 | Tps_LRv5b_scf10 | 8.00E-06 |
| Female | Tps_LRv5b_scf8 | Tps_LRv5b_scf11 | 0.013 |
| Female | Tps_LRv5b_scf8 | Tps_LRv5b_scf12 | 0.53 |
| Female | Tps_LRv5b_scf9 | Tps_LRv5b_scf10 | 0.29 |
| Female | Tps_LRv5b_scf9 | Tps_LRv5b_scf11 | 6.80E-08 |
| Female | Tps_LRv5b_scf9 | Tps_LRv5b_scf12 | 0.07 |
| Female | Tps_LRv5b_scf10 | Tps_LRv5b_scf11 | 1.90E-12 |
| Female | Tps_LRv5b_scf10 | Tps_LRv5b_scf12 | 0.0014 |
| Female | Tps_LRv5b_scf11 | Tps_LRv5b_scf12 | 0.0035 |
| Male | Tps_LRv5b_scf1 | Tps_LRv5b_scf2 | 0.03 |
| Male | Tps_LRv5b_scf1 | Tps_LRv5b_scf3 | 1.80E-18 |
| Male | Tps_LRv5b_scf1 | Tps_LRv5b_scf4 | 0.0022 |
| Male | Tps_LRv5b_scf1 | Tps_LRv5b_scf5 | 6.40E-05 |
| Male | Tps_LRv5b_scf1 | Tps_LRv5b_scf6 | 0.28 |
| Male | Tps_LRv5b_scf1 | Tps_LRv5b_scf7 | 0.97 |
| Male | Tps_LRv5b_scf1 | Tps_LRv5b_scf8 | 0.0033 |
| Male | Tps_LRv5b_scf1 | Tps_LRv5b_scf9 | 0.35 |
| Male | Tps_LRv5b_scf1 | Tps_LRv5b_scf10 | 3.40E-05 |
| Male | Tps_LRv5b_scf1 | Tps_LRv5b_scf11 | 2.60E-08 |
| Male | Tps_LRv5b_scf1 | Tps_LRv5b_scf12 | 0.16 |
| Male | Tps_LRv5b_scf2 | Tps_LRv5b_scf3 | 1.40E-09 |
| Male | Tps_LRv5b_scf2 | Tps_LRv5b_scf4 | 0.35 |
| Male | Tps_LRv5b_scf2 | Tps_LRv5b_scf5 | 0.065 |
| Male | Tps_LRv5b_scf2 | Tps_LRv5b_scf6 | 0.57 |
| Male | Tps_LRv5b_scf2 | Tps_LRv5b_scf7 | 0.1 |
| Male | Tps_LRv5b_scf2 | Tps_LRv5b_scf8 | 0.28 |
| Male | Tps_LRv5b_scf2 | Tps_LRv5b_scf9 | 0.018 |
| Male | Tps_LRv5b_scf2 | Tps_LRv5b_scf10 | 7.60E-08 |
| Male | Tps_LRv5b_scf2 | Tps_LRv5b_scf11 | 0.00065 |
| Male | Tps_LRv5b_scf2 | Tps_LRv5b_scf12 | 0.97 |
| Male | Tps_LRv5b_scf3 | Tps_LRv5b_scf4 | 3.40E-06 |
| Male | Tps_LRv5b_scf3 | Tps_LRv5b_scf5 | 4.00E-04 |
| Male | Tps_LRv5b_scf3 | Tps_LRv5b_scf6 | 5.60E-09 |
| Male | Tps_LRv5b_scf3 | Tps_LRv5b_scf7 | 3.50E-11 |
| Male | Tps_LRv5b_scf3 | Tps_LRv5b_scf8 | 8.50E-05 |
| Male | Tps_LRv5b_scf3 | Tps_LRv5b_scf9 | 1.90E-11 |
| Male | Tps_LRv5b_scf3 | Tps_LRv5b_scf10 | 1.60E-23 |
| Male | Tps_LRv5b_scf3 | Tps_LRv5b_scf11 | 0.081 |
| Male | Tps_LRv5b_scf3 | Tps_LRv5b_scf12 | 1.30E-05 |
| Male | Tps_LRv5b_scf4 | Tps_LRv5b_scf5 | 0.39 |
| Male | Tps_LRv5b_scf4 | Tps_LRv5b_scf6 | 0.19 |
| Male | Tps_LRv5b_scf4 | Tps_LRv5b_scf7 | 0.02 |
| Male | Tps_LRv5b_scf4 | Tps_LRv5b_scf8 | 0.81 |
| Male | Tps_LRv5b_scf4 | Tps_LRv5b_scf9 | 0.0037 |
| Male | Tps_LRv5b_scf4 | Tps_LRv5b_scf10 | 6.20E-09 |
| Male | Tps_LRv5b_scf4 | Tps_LRv5b_scf11 | 0.019 |
| Male | Tps_LRv5b_scf4 | Tps_LRv5b_scf12 | 0.53 |
| Male | Tps_LRv5b_scf5 | Tps_LRv5b_scf6 | 0.033 |
| Male | Tps_LRv5b_scf5 | Tps_LRv5b_scf7 | 0.0021 |
| Male | Tps_LRv5b_scf5 | Tps_LRv5b_scf8 | 0.6 |
| Male | Tps_LRv5b_scf5 | Tps_LRv5b_scf9 | 0.00037 |
| Male | Tps_LRv5b_scf5 | Tps_LRv5b_scf10 | 1.30E-10 |
| Male | Tps_LRv5b_scf5 | Tps_LRv5b_scf11 | 0.17 |
| Male | Tps_LRv5b_scf5 | Tps_LRv5b_scf12 | 0.21 |
| Male | Tps_LRv5b_scf6 | Tps_LRv5b_scf7 | 0.34 |
| Male | Tps_LRv5b_scf6 | Tps_LRv5b_scf8 | 0.15 |
| Male | Tps_LRv5b_scf6 | Tps_LRv5b_scf9 | 0.1 |
| Male | Tps_LRv5b_scf6 | Tps_LRv5b_scf10 | 1.90E-05 |
| Male | Tps_LRv5b_scf6 | Tps_LRv5b_scf11 | 0.00048 |
| Male | Tps_LRv5b_scf6 | Tps_LRv5b_scf12 | 0.66 |
| Male | Tps_LRv5b_scf7 | Tps_LRv5b_scf8 | 0.018 |
| Male | Tps_LRv5b_scf7 | Tps_LRv5b_scf9 | 0.5 |
| Male | Tps_LRv5b_scf7 | Tps_LRv5b_scf10 | 0.0015 |
| Male | Tps_LRv5b_scf7 | Tps_LRv5b_scf11 | 1.10E-05 |
| Male | Tps_LRv5b_scf7 | Tps_LRv5b_scf12 | 0.27 |
| Male | Tps_LRv5b_scf8 | Tps_LRv5b_scf9 | 0.0031 |
| Male | Tps_LRv5b_scf8 | Tps_LRv5b_scf10 | 2.80E-08 |
| Male | Tps_LRv5b_scf8 | Tps_LRv5b_scf11 | 0.068 |
| Male | Tps_LRv5b_scf8 | Tps_LRv5b_scf12 | 0.46 |
| Male | Tps_LRv5b_scf9 | Tps_LRv5b_scf10 | 0.033 |
| Male | Tps_LRv5b_scf9 | Tps_LRv5b_scf11 | 2.30E-06 |
| Male | Tps_LRv5b_scf9 | Tps_LRv5b_scf12 | 0.1 |
| Male | Tps_LRv5b_scf10 | Tps_LRv5b_scf11 | 5.70E-14 |
| Male | Tps_LRv5b_scf10 | Tps_LRv5b_scf12 | 6.30E-05 |
| Male | Tps_LRv5b_scf11 | Tps_LRv5b_scf12 | 0.013 |

**Supplemental Figures**

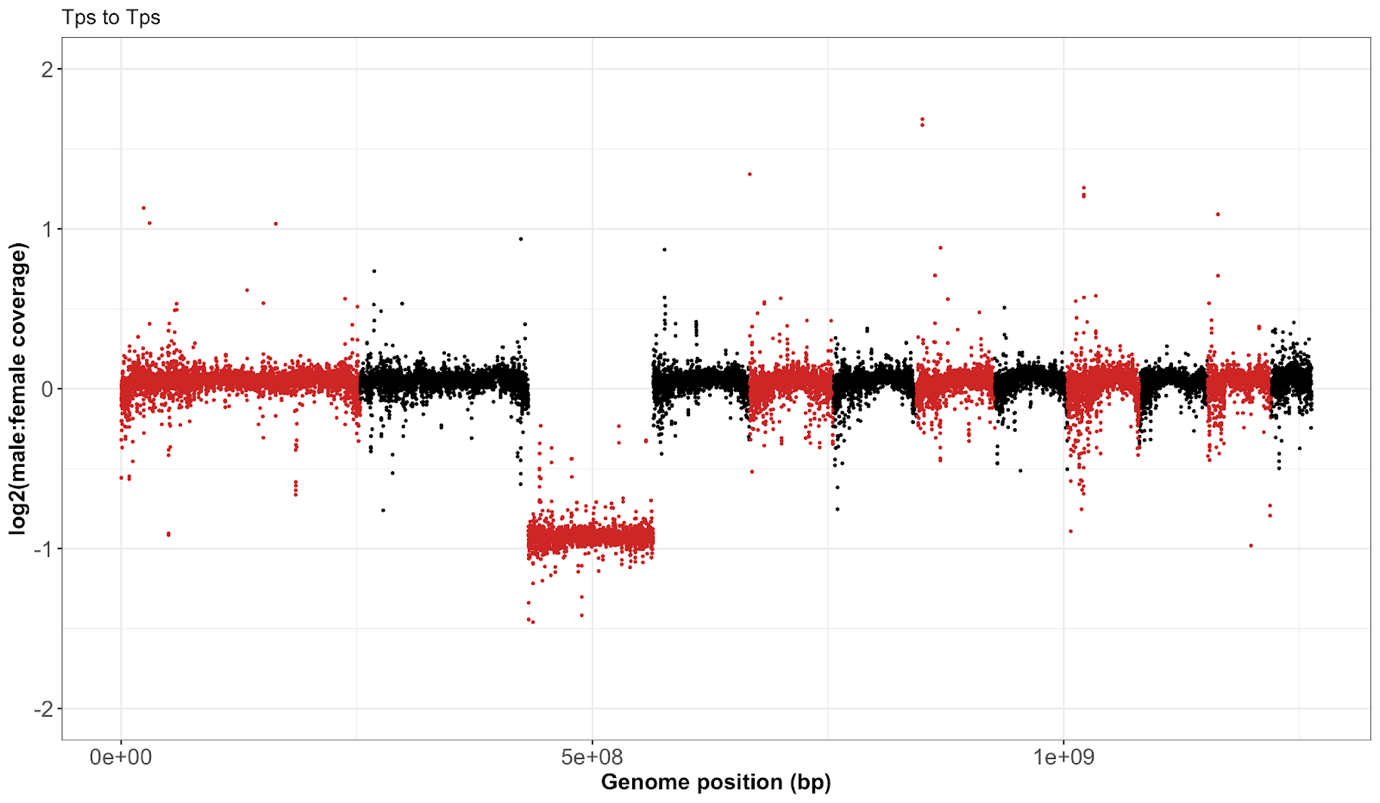

**Supplemental Figure S1. *T. poppense* X chromosome identification.** The plot shows the log_2_ ratio of male to female coverage of 100 kb sliding windows across the genome. Alternated colours designate different chromosomes, with chromosome 3 showing a much lower overall male to female coverage ratio.

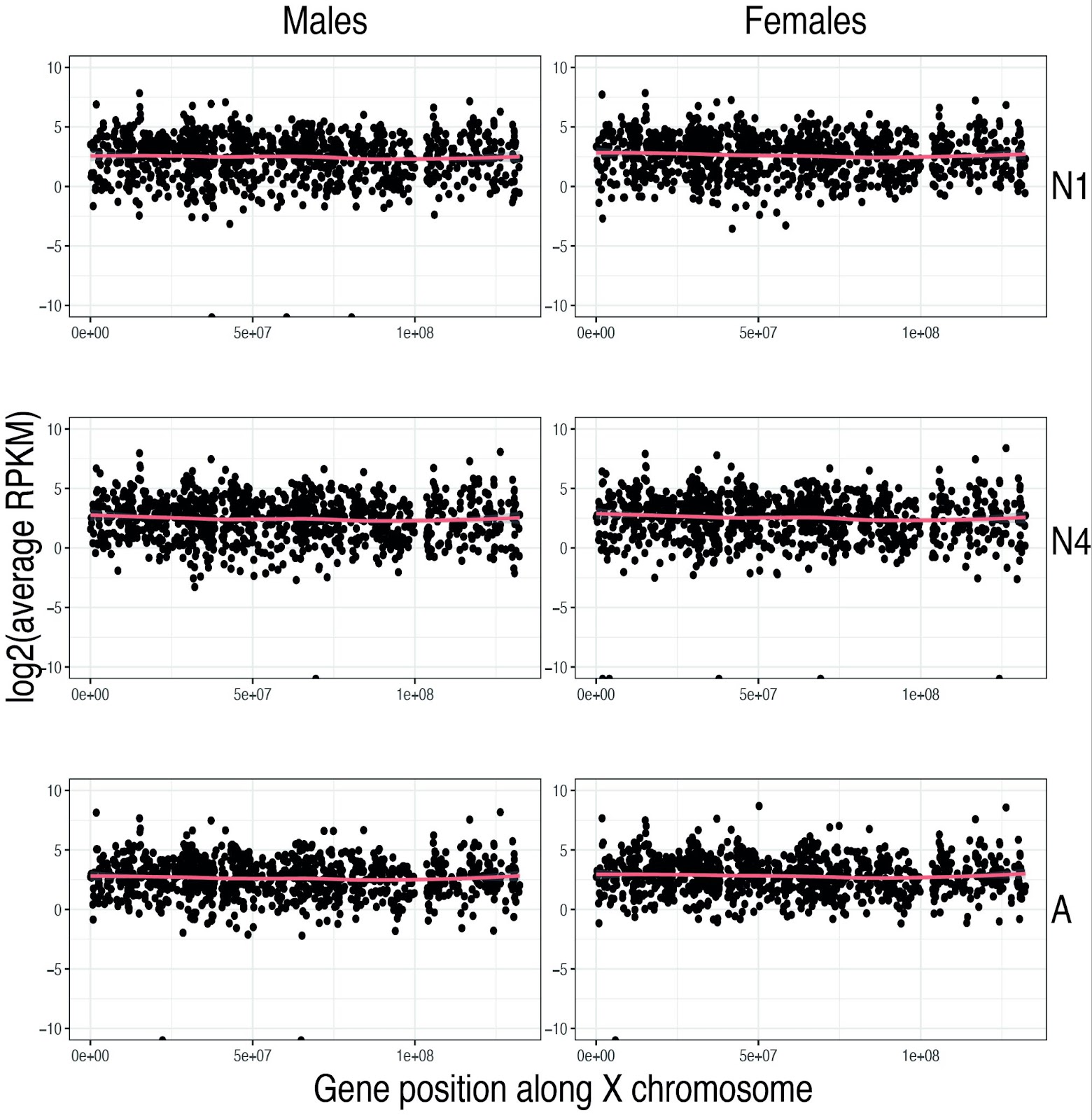

**Supplemental Figure S2.** Mean expression levels per gene along the X chromosome in brain somatic tissue at 1st (N1), 4th (N4) and adult (A) stages in females (left) and males (right). The line in each panel represents a loess smoothed curve.

 
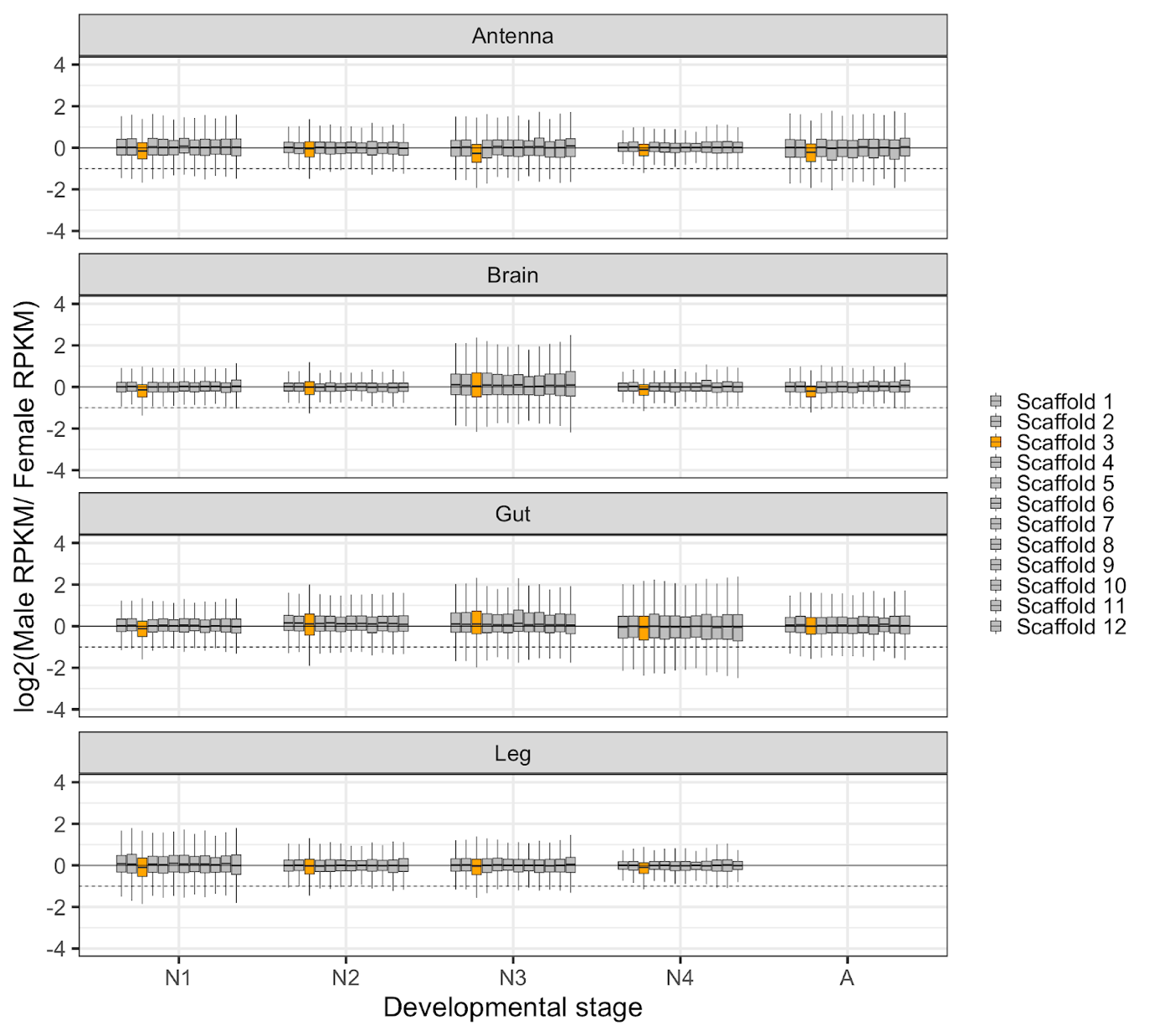

**Supplemental Figure S3.** Log2 of male to female expression ratio across scaffolds (representing all 12 *T. poppense* chromosomes) at five developmental stages: N1, N2, N3, N4, and A (representing the 1st to 4th nymphal stages and the adult stage, respectively). Scaffold 3, represented in orange, corresponds to the X chromosome, while autosomal scaffolds are depicted in gray. The panels, arranged from top to bottom, showcase the Log2 ratio in different somatic tissues: antenna, brain, guts, and legs. Boxplots depict the median, the lower and upper quartiles, while the whiskers represent the minimum and maximum values, within 1.5x the interquartile range.

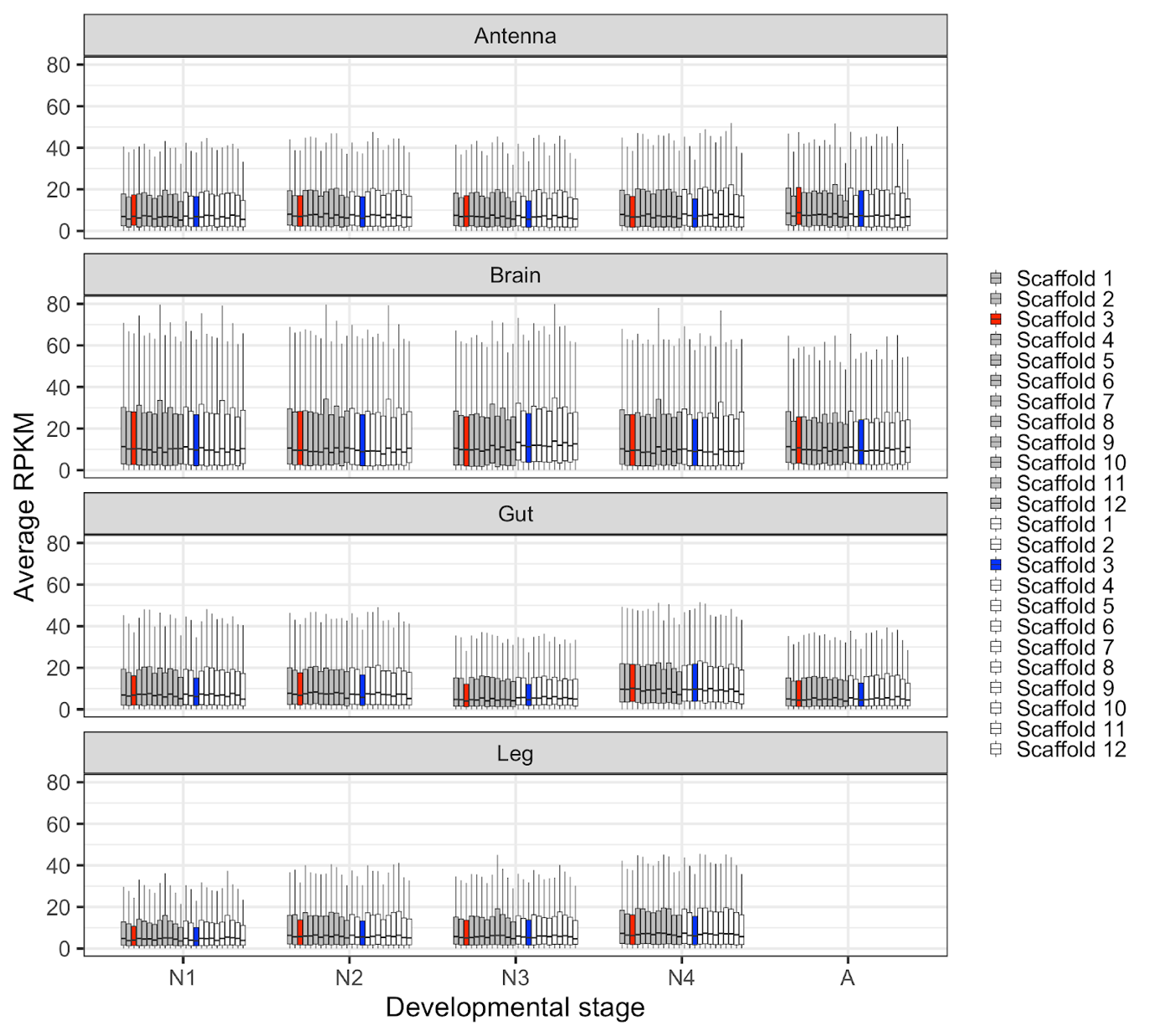

**Supplemental Figure S4.** Average RPKM expression levels across scaffolds in females (grey) and in males (white) along development, scaffold three corresponds to the X and is depicted in red (females) and blue (males). The panels, from top to bottom, showcase the Average RPKM in different somatic tissues: antenna, brain, guts, and legs. Boxplots depict the median, the lower and upper quartiles, while the whiskers represent the minimum and maximum values, within 1.5x the interquartile range.

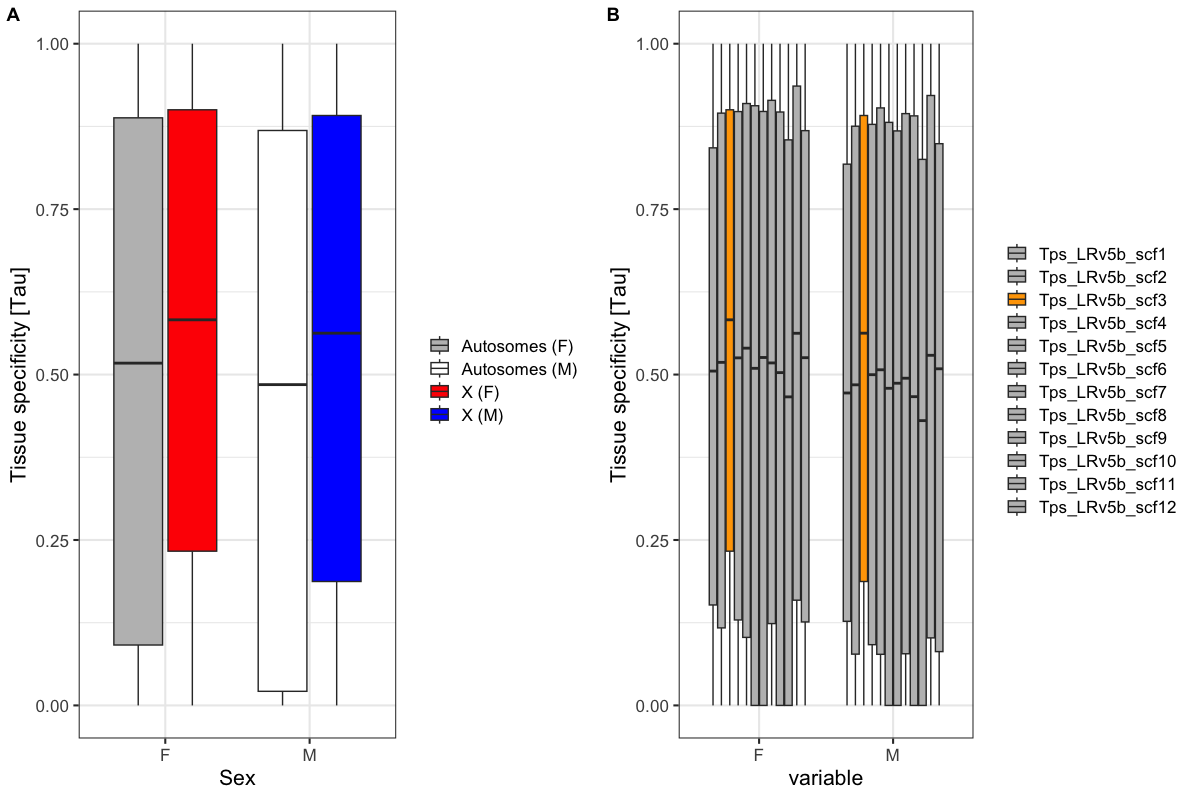

**Supplemental Figure S5. A)** Tissue specificity of X chromosomes in males (M) (blue box) and females (F) (red box) as compared to the autosomes (white and gray boxes), calculated across three somatic tissues. Because genes on the X are often testes or ovaries specific, we here repeated the analysis presented in the main text based on four tissues (three somatic tissues and reproductive tracts) with the three somatic tissues only, and X tissue specificity remained higher than autosomes Wilcoxon test, *p*_adj (females)_= 6.7e-13, *p*_adj (males)_= 2.4e-16 **B)** Tissue specificity in females (F) and males (M) across scaffolds (see Supplemental Table 5), based on three somatic tissues. Scaffold 3, represented in orange, corresponds to the X chromosome, while other scaffolds are depicted in gray. Boxplots depict the median, the lower and upper quartiles, while the whiskers represent the minimum and maximum values, within 1.5x the interquartile range.

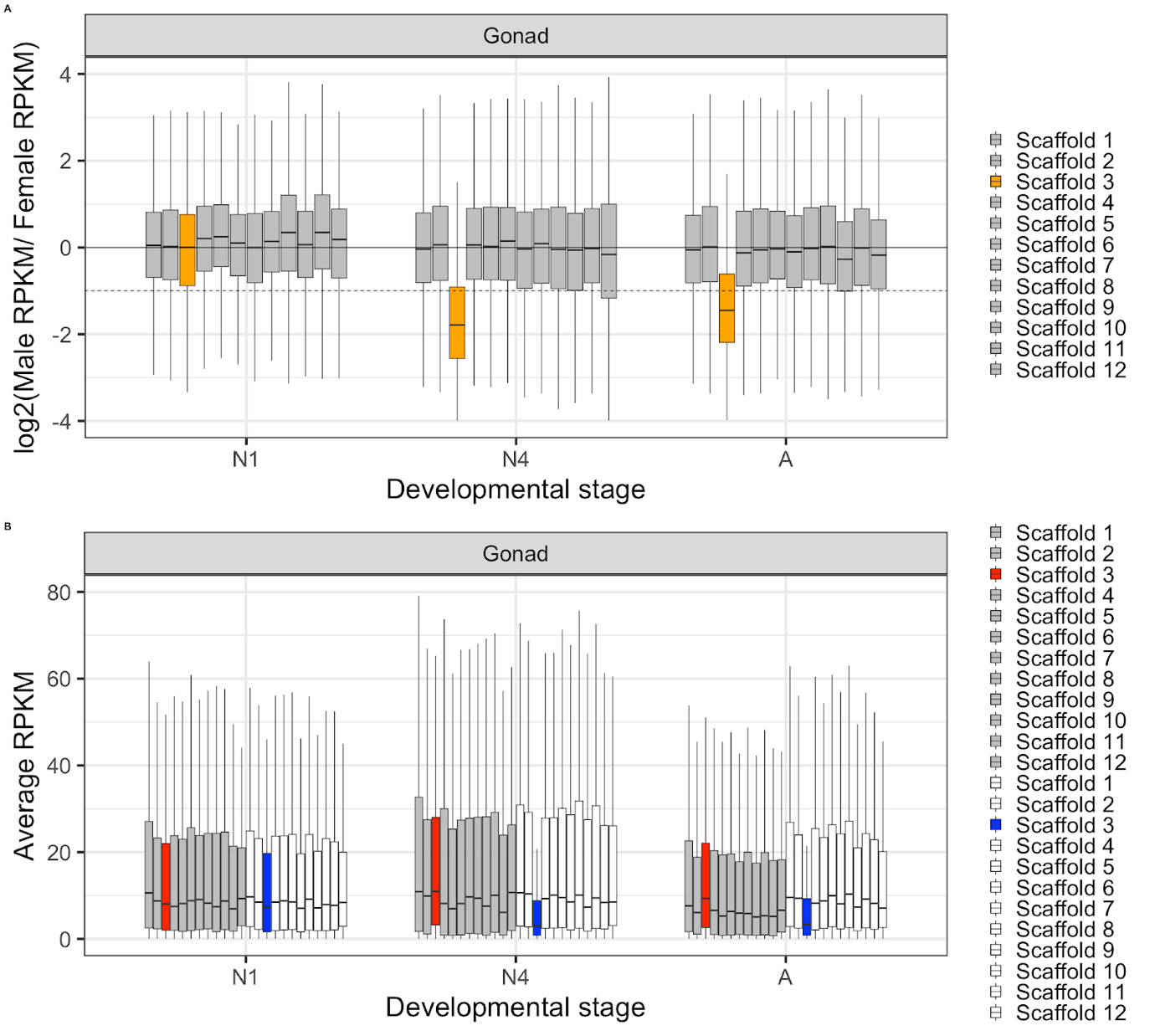

**Supplemental Figure S6.** **A)** The Log2 ratio of RPKM levels of expression between males and females across scaffolds at three developmental stages (N1, N4, and Adult) in the reproductive tract. Scaffold 3, represented in orange, corresponds to the X chromosome, while other scaffolds are depicted in gray. **B)**  Average RPKM expression levels at three developmental stages (N1, N4, and adult) in the reproductive tract separated for different scaffolds (chromosomes) in females (grey) and in males (white) boxes, scaffold three corresponds to the X and is depicted in red (females) and blue (males). Boxplots depict the median, the lower and upper quartiles, while the whiskers represent the minimum and maximum values, within 1.5x the interquartile range.

 
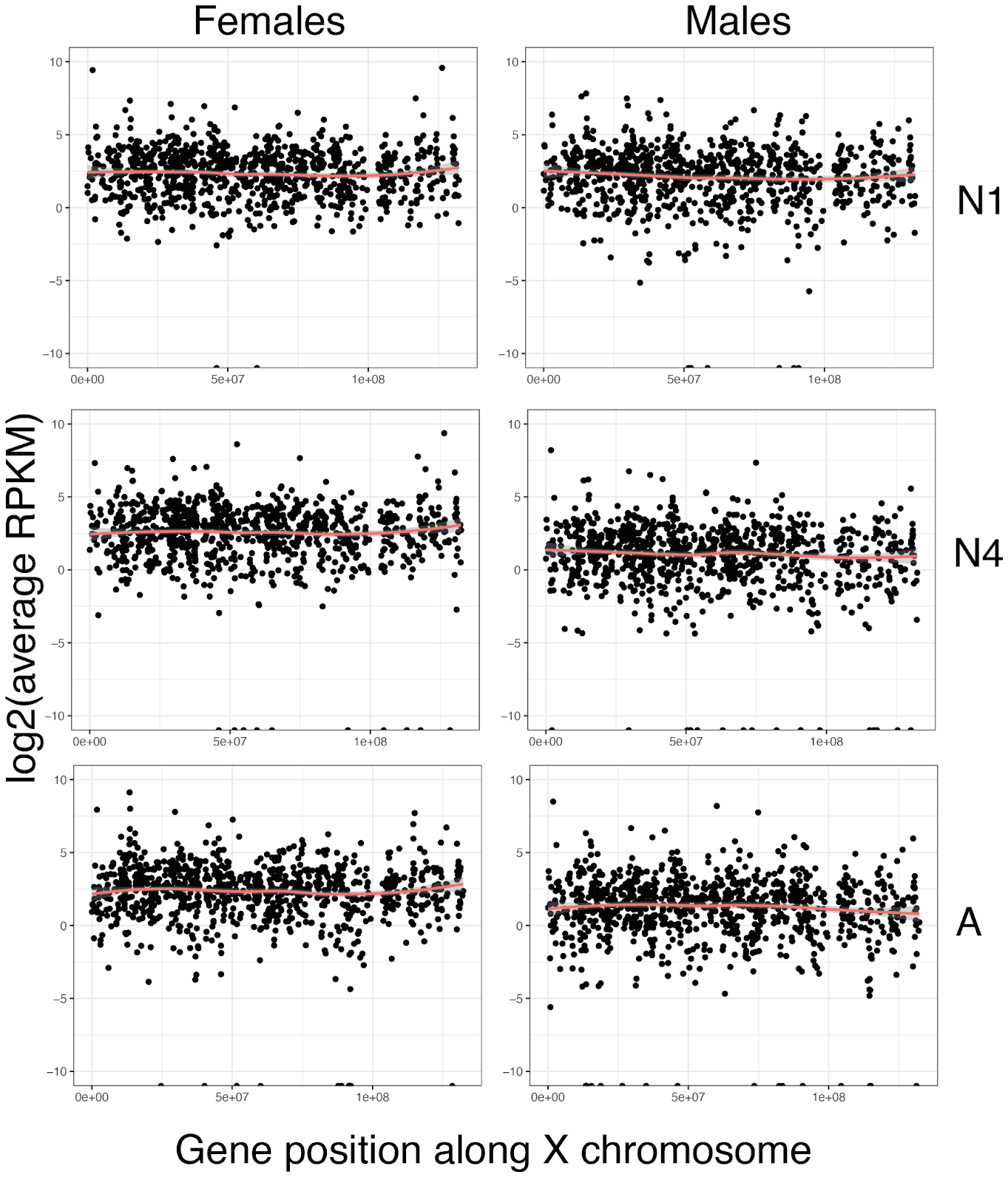

**Supplemental Figure S7.** Mean expression RPKM levels per gene along the X chromosome at 1^st^ (N1), 4^th^ (N4) and Adult (A) stages in reproductive tracts of females (left) and males (right) panels. The red line in each panel shows the loess smoothed curve.

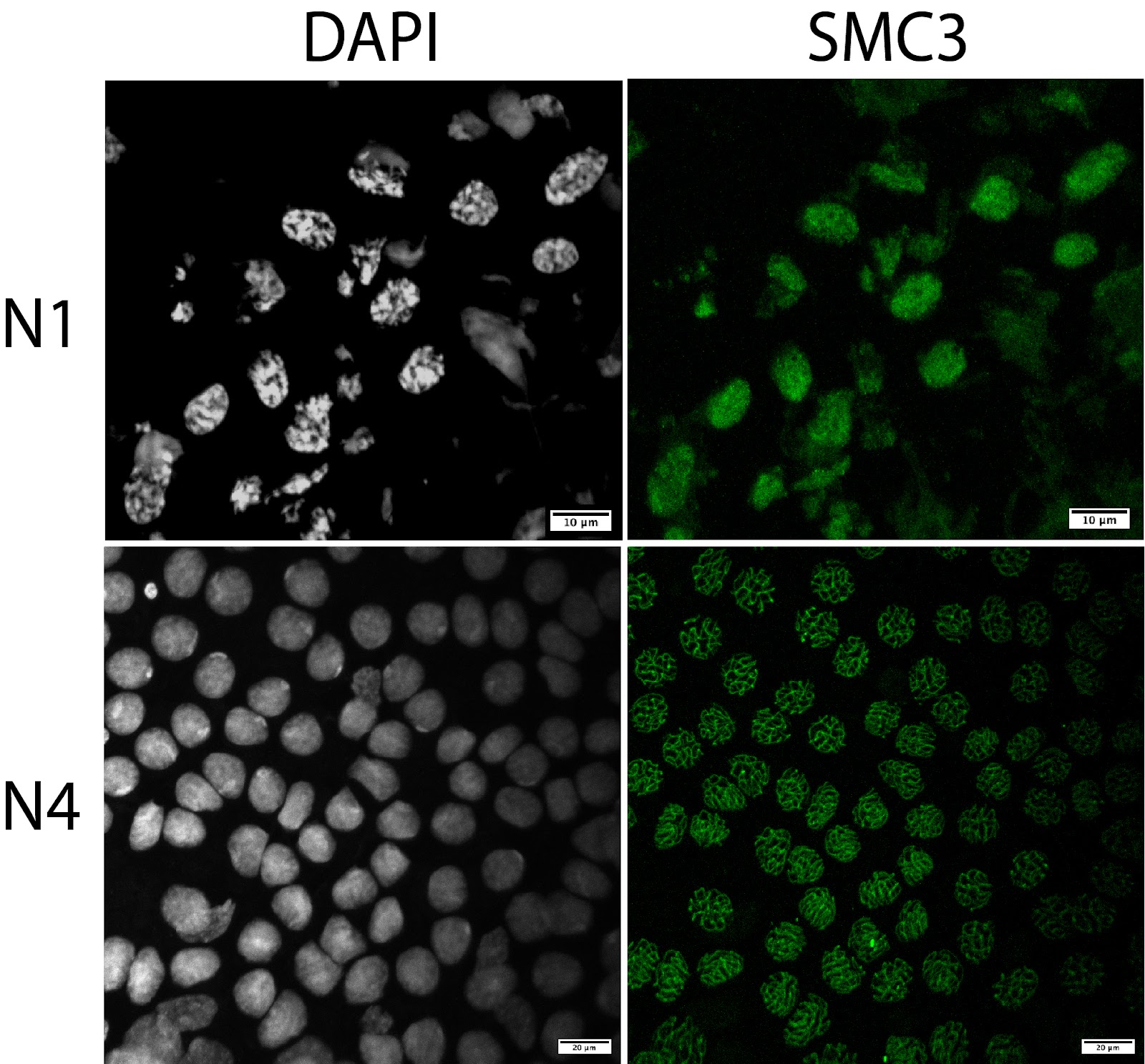

**Supplemental Figure S8**. DAPI, SMC3 staining applied to testes squashes of 1^st^ (N1) and 4^th^ (N4) nymphal stage males. SMC3 is a protein of the cohesin complex marking chromosome axes during meiosis 1. No cells in meiosis I were detected at N1, while numerous cells at this stage were detected at N4.
